# Differential turnover of Nup188 controls its levels at centrosomes and role in centriole duplication

**DOI:** 10.1101/664409

**Authors:** Nidhi Vishnoi, Karthigeyan Dhanasekeran, Madeleine Chalfant, Ivan Surovstev, Mustafa K. Khokha, C. Patrick Lusk

## Abstract

*NUP188* encodes a scaffold component of the nuclear pore complex (NPC) and has been implicated as a congenital heart disease gene through an ill-defined function at centrioles. Here, we explore the mechanisms that physically and functionally segregate Nup188 between the pericentriolar material (PCM) and NPCs throughout the cell cycle. Pulse-chase fluorescent labeling approaches indicate that Nup188 populates centrosomes with newly synthesized protein that does not exchange with NPCs even after mitotic NPC breakdown. In addition, the steady-state level of Nup188 at centrosomes is controlled by the sensitivity of the PCM pool, but not the NPC pool, to proteasomal degradation. Proximity-labeling and super-resolution microscopy supports that Nup188 interacts with components of PCM including Cep192 and the centriolar satellite component, PCM1. Consistent with this, Nup188 plays a role in centriole duplication at or upstream of Sas6 loading. Together, our data establish Nup188 as a functional component of PCM and potentially provides insight into the pathogenesis of congenital heart disease.

## Introduction

The enclosure of the genome within the nuclear membranes occurred alongside the evolution of nuclear pore complexes (NPCs), which control all molecular traffic between the nucleus and cytoplasm. There are ∼30 nucleoporins or “nups” that, alongside nup binding partners, construct modular subcomplex building blocks that come together in multiples of eight to assemble ∼100 MD transport channels (Hampoelz et al., 2019). The major architectural units of the NPC scaffold consist of the Nup107-160, “Y” or “outer ring” complex in addition to the Nup93 or “inner ring complex” (Amlacher et al., 2011; Bui et al., 2013; Kim et al., 2018; Kosinski et al., 2016; Siniossoglou et al., 2000; von Appen et al., 2015). The latter consists of Nup93, Nup155, Nup35 (Nup53), Nup205 and Nup188 (Amlacher et al., 2011; Vollmer and Antonin, 2014). The ring complexes provide anchor points for Phe-Gly (FG)-rich nups that establish a size-selective diffusion barrier and provide binding sites for shuttling nuclear transport receptors (NTRs/karyopherins/importins/exportins) bound to cargo (Schmidt and Gorlich, 2016; Wente and Rout, 2010).

In addition to their well-established roles at NPCs, some nups moonlight in other subcellular locations like in the nucleus (Capelson et al., 2010; Capitanio et al., 2018; Kalverda et al., 2010; Liang et al., 2013; Vaquerizas et al., 2010), or by binding the mitotic apparatus (Wozniak et al., 2010). For example, a fraction of the Nup107-160 complex is recruited to kinetochores after nuclear envelope and NPC breakdown during mitosis (Belgareh et al., 2001; Loiodice et al., 2004; Zuccolo et al., 2007) where it helps to recruit the γ-Tubulin ring complex (Mishra et al., 2010). This association might also recruit NTRs and components of the Ran GTPase system, which also play a central role in spindle assembly (Clarke and Zhang, 2008; Zhang et al., 2014). Other nups have also been shown to interact with the mitotic spindle (Cross and Powers, 2011) and spindle assembly checkpoint components (Iouk et al., 2002; Lussi et al., 2010; Markossian et al., 2015; Rodenas et al., 2012; Rodriguez-Bravo et al., 2014; Schweizer et al., 2013). Further, there is evidence to support that both Nup62 (Hashizume et al., 2013) and Nup188 (Itoh et al., 2013) localize to centrosomes, the major microtubule organizing centers in mammalian cells. In general, the molecular mechanisms that define nup function in association with the mitotic apparatus remain to be fully defined.

Understanding the full spectrum of nup function is becoming more pressing as increasing evidence supports that disruption of the nuclear transport system is causative of a wide range of neurodegenerative diseases (Sakuma and D’Angelo, 2017) and cancers (Kohler and Hurt, 2010; Rodriguez-Bravo et al., 2018; Simon and Rout, 2014). In addition, modern patient genomics is revealing a remarkable list of nup disease variants associated with, for example, Triple A syndrome (Tullio-Pelet et al., 2000), steroid resistant nephrotic syndrome (Braun et al., 2018; Braun et al., 2016; Miyake et al., 2015), non-progressive congenital ataxia (Zanni et al., 2019) and heterotaxy (Fakhro et al., 2011; Manheimer et al., 2018). Heterotaxy is a disorder of left-right patterning that can lead to mispositioned hearts and a severe form of congenital heart disease (Sutherland and Ware, 2009). Interestingly, other nups like Gle1 have also been associated with left-right patterning in Zebrafish suggesting a specific role for nups in development (Jao et al., 2017; Kaneb et al., 2015). Indeed, while it is likely that some disease-causing nup malfunction is linked directly to defects in nuclear transport, our prior investigation of a copy number variant of *NUP188* in a heterotaxy patient (Fakhro et al., 2011) suggested a role for Nup188 at the bases of cilia, key organelles essential for left-right patterning in the developing embryo (Del Viso et al., 2016).

Cilia are built atop centrioles, which are ancient organelles that consist of 9-fold radially-arranged triplets of microtubules (Azimzadeh and Marshall, 2010; Gonczy, 2012; Nigg and Holland, 2018). While their most evolutionarily conserved function is thought to be the formation of cilia, centrioles also form the core of centrosomes by acquiring pericentriolar material (PCM) consisting of γ-Tubulin ring complexes (γ-TURCs) and centrosome proteins (Ceps)(among many others)(Andersen et al., 2003; Jakobsen et al., 2011). During interphase, PCM is organized into multiple distinct “layers” with unique molecular components in each (Fry et al., 2017; Lawo et al., 2012; Mennella et al., 2012; Sonnen et al., 2012). For example, Cep192 is a major component of the inner layer, with Cep152 and Pericentrin each populating the intermediate and outer layers, respectively (Lawo et al., 2012; Mennella et al., 2012; Sonnen et al., 2012). Interestingly, this distinct radial-layer architecture is altered as cells enter G2/M when centrosomes “mature” with the addition of more PCM (Haren et al., 2009; Khodjakov and Rieder, 1999; Lawo et al., 2012; Lee and Rhee, 2011; Sonnen et al., 2012; Zhu et al., 2008). In addition, the delivery of PCM components to the centrosome is thought to require, in some cases, an intermediary in the form of centriolar satellites, cytosolic granules that have PCM1 as their foundational molecular component (Gupta et al., 2015; Hori and Toda, 2017; Kubo et al., 1999; Tollenaere et al., 2015). Mature centrosomes contribute to spindle formation and the successful capture and segregation of chromosomes (Prosser and Pelletier, 2017).

After chromosome segregation, daughter cells inherit a single centrosome consisting of two centrioles, only one of which has PCM; the other acquires PCM through a centriole-to-centrosome conversion process in G1 (Fong et al., 2018; Fu et al., 2016; Izquierdo et al., 2014; Kim et al., 2019). To double centrosome number, each of the centrioles must first be duplicated in S-phase by a nascent “pro” centriole that assembles perpendicularly from the parent centriole outer wall (Nigg and Holland, 2018). This is achieved by a highly coordinated series of yet to be fully defined molecular events triggered by the recruitment of polo-like-kinase 4 (Plk4)(Bettencourt-Dias et al., 2005; Habedanck et al., 2005; O’Connell et al., 2001) by Cep152 and Cep192 (Cizmecioglu et al., 2010; Hatch et al., 2010; Kim et al., 2013; Park et al., 2014; Sonnen et al., 2013) and its phosphorylation of a network of proteins including STIL, CPAP and Sas6 (Dammermann et al., 2004; Delattre et al., 2004; Dzhindzhev et al., 2017; Kemp et al., 2004; Kirkham et al., 2003; Kratz et al., 2015; Leidel et al., 2005; Leidel and Gonczy, 2003; Moyer et al., 2015; Moyer and Holland, 2019; Ohta et al., 2014; Pelletier et al., 2004; Stevens et al., 2010; Tang et al., 2011). Sas6 helps form a supramolecular cartwheel-shaped structure that serves as a scaffold for microtubule nucleation (Gonczy, 2012). Procentrioles acquire additional proteins that help control microtubule growth (Nigg and Holland, 2018), however, they are not considered to be fully mature centrioles until they have acquired distal and subdistal appendages, and with them, the capability to form cilia (Hoyer-Fender, 2010).

Here, we build on our prior work suggesting that Nup188 surrounds the centrioles at cilia bases (Del Viso et al., 2016). We specifically explore the mechanisms that lead to Nup188 recruitment to PCM and determine that newly synthesized Nup188 populates PCM at all stages of the cell cycle. Our data support a model where modulating Nup188 turnover rates ultimately controls its steady state levels at centrosomes. We further uncover physical interactions with established PCM components and provide evidence that Nup188 is required for centriole duplication. Thus, Nup188 is a shared component of both NPCs and PCM required for the centriole life cycle.

## Results

### Nup188 accumulates at centrosomes with cyclin-like behavior

To better determine the mechanism of Nup188 recruitment to PCM, we first examined the steady-state distribution of Nup188 in interphase and mitosis. Immunofluorescence microscopy showed *α*-Nup188 staining at the nuclear periphery and in the nucleus of interphase cells and a spatially distinct pool that colocalized with γ-Tubulin, which labels centrosomes (*α*-γ-Tubulin, red)(Figure 1A). Interestingly, we observed a continuous increase of *α*-Nup188 signal at centrosomes as cells progressed into mitosis, reaching a peak in metaphase with ∼3 fold higher levels of fluorescence relative to interphase levels (mitosis stage assessed with Hoechst staining) before diminishing in telophase (Figure 1A, B). These results are consistent with published work (Itoh et al., 2013), and the temporal signature of this accumulation (and loss) is similar to mitotic cyclins and to other PCM components that drive centrosome maturation during mitosis, including γ-Tubulin (Figure 1C) (Woodruff et al., 2014).

**Figure 1.**
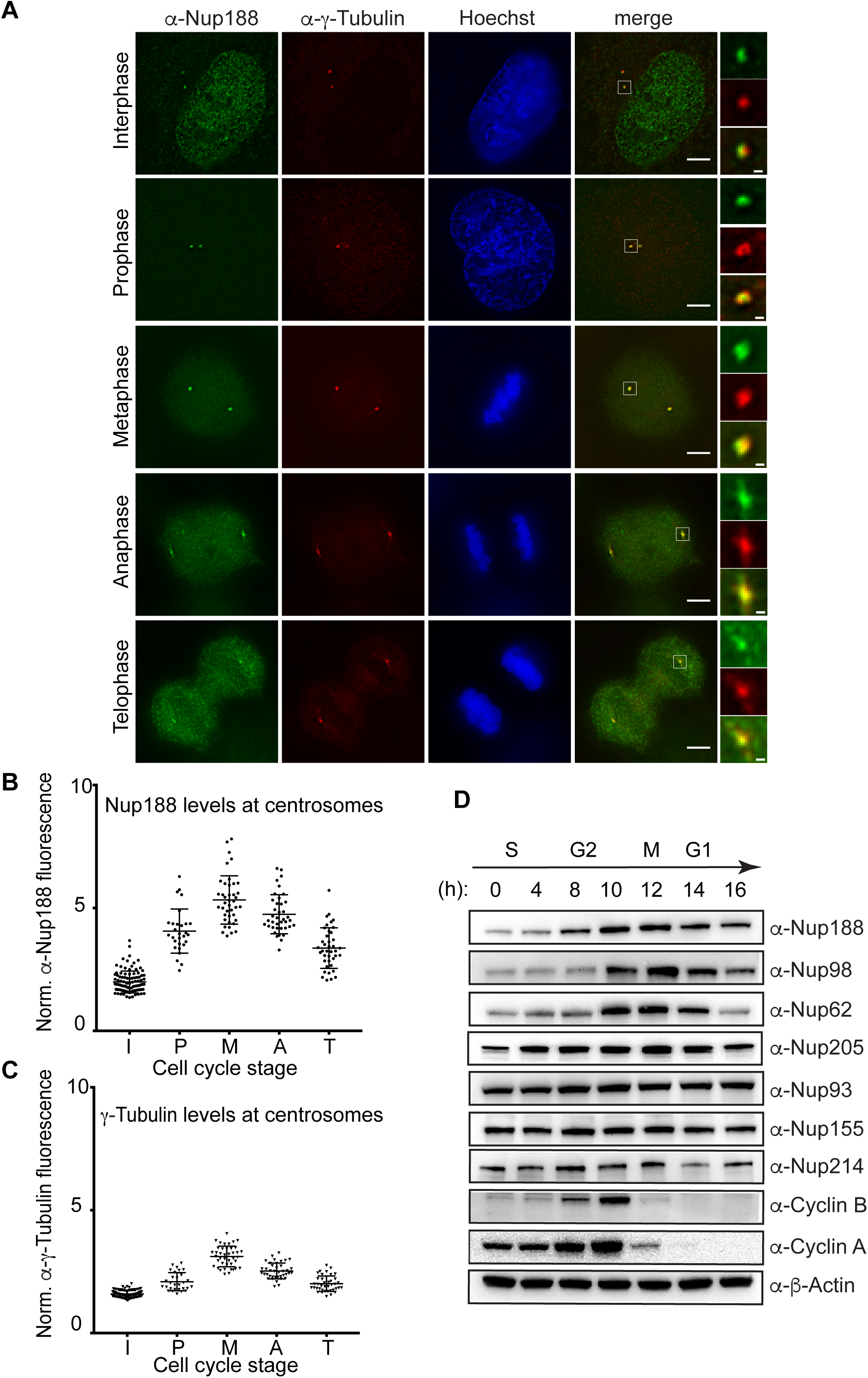
Nup188 levels increase at centrosomes during mitosis. **(A)** Immunofluorescence micrographs of HeLa-M cells stained with *α*-Nup188 (green) and *α*-γ-Tubulin (red) antibodies (with merge) during interphase and mitosis. Mitotic stages were defined by the morphology of the Hoechst-stained chromosomes (blue). Scale bar is 5 μm. Magnifications of the boxed area encompassing centrosomes (green, red and merge) are shown at right. Scale bar is 0.5 μm. **(B and C)** Plot of the total fluorescence (in arbitrary units) of *α*-Nup188 (B) and *α*-γ-Tubulin (C) staining fluorescence of individual centrosomes normalized (norm.) to background (extracellular) fluorescence in interphase (I) (n=100), prophase (P) (n=29), metaphase (M) (n=41), anaphase (A) (n=39) and telophase (T) (n=40) from three independent experiments. Mean ±SD are indicated. **(D)** Several nups including Nup188 exhibit cyclin-like behavior. Western blots assessing the indicated nup levels in total protein samples generated from HeLa-M cells synchronized and released from S-phase. Approximate cell cycle stage is indicated at top and is informed by assessing Cyclin A and B synthesis and degradation. *α*-β-Actin is used to assess total protein load in each sample.

We wondered whether this increase in *α*-Nup188 staining reflected a rise in total levels of Nup188 during mitosis. We therefore arrested cells in S-phase and synchronously released them into the cell cycle while monitoring nup levels by Western blot (Figure 1D). To monitor synchronous entry into mitosis, the increase and rapid clearance of Cyclin A and Cyclin B, which demarcate the G2-M and the metaphase-anaphase transitions, respectively (Furuno et al., 1999; Pines and Hunter, 1989) was simultaneously assessed (Figure 1D). Nup188 levels began rising at 8 h post S-phase release and peaked at 12 h, which corresponds to a timepoint just after Cyclin B degradation. Interestingly, this behavior is not mirrored by other nups like Nup214 and the other components of the inner ring complex including Nup93 and Nup155, whose levels remain roughly consistent through the timecourse; although, we observed a modest increase for Nup205, the paralog of Nup188. We next assessed additional nups that have been functionally linked to cell cycle progression including the FG-nups, Nup62 (Hashizume et al., 2013; Wu et al., 2016) and Nup98 (Cross and Powers, 2011). Here, we observed an even more striking cyclin-like behavior: both of these proteins increased and then rapidly diminished with kinetics that mirrored those of Cyclin B. We further note that Nup96, but not other components of the outer ring complex, was previously shown to decline in mitosis as well (Chakraborty et al., 2008). Together, these correlations between nup levels and cell cycle stage support that some nups, including Nup188, respond to cell cycle cues.

### Centrosomal Nup188 is populated by newly synthesized protein, not from NPCs

We next asked whether the increase of Nup188 at centrosomes during mitosis was derived from NPC breakdown, or, from new protein synthesis. To test this, we generated a stable HeLa cell line that expresses a SNAP-tagged Nup188 fusion protein behind a doxycycline (dox)-inducible promoter. To reduce the potential for overexpression artifacts, we established conditions where SNAP-Nup188 is produced below endogenous levels (i.e. undetectable by the *α*-Nup188 antibody, Figure S1A). We also ensured that SNAP-Nup188 showed cyclin-like behavior just like the endogenous protein (Figure 2A).

**Figure 2.**
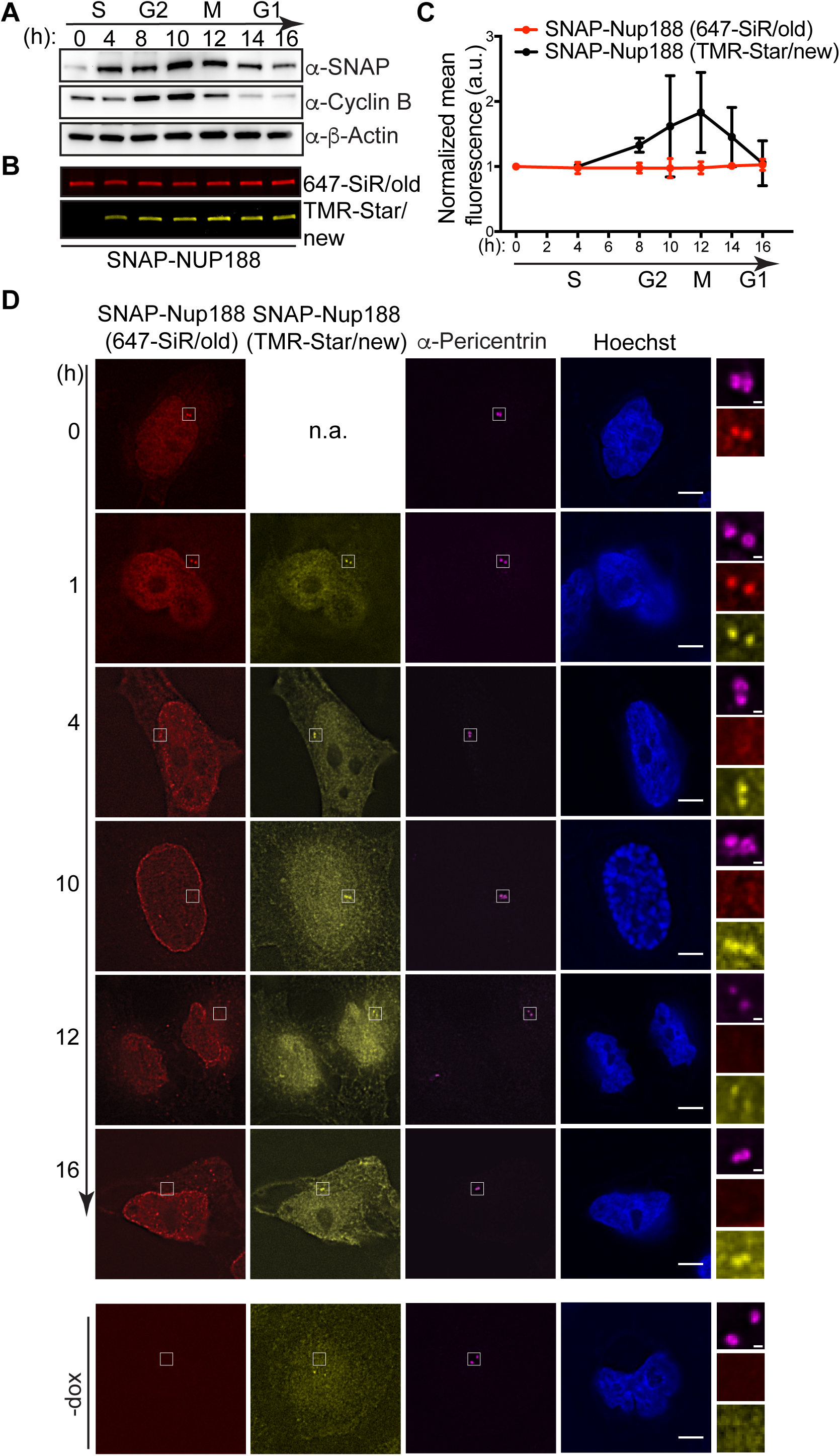
Centrosomes are populated by newly synthesized Nup188 not exchange from NPCs. **(A)** Like endogenous Nup188, SNAP-Nup188 undergoes mitotic oscillation. Western blots examining the levels of SNAP-Nup188 in total protein samples derived from synchronized cells. Approximate cell cycle stage is indicated at top referencing Cyclin B synthesis and degradation; *α*-β-Actin is used to assess total protein loads. **(B)** As in A, but SNAP-Nup188 protein is visualized by fluorescence. SNAP-Nup188 is first pulse-labeled with a 647-SiR-dye (red/”old”) and then chase-labeled at the indicated time points with a TMR-Star dye (yellow/”new”). **(C)** Plot of fluorescence from (B) with two additional experimental replicates. The mean fluorescence from these three experiments is represented ±SD. **(D)** Fluorescence images of HeLa cells expressing SNAP-Nup188 were synchronized with thymidine at S-phase and labeled with 647-SiR dye (red). Subsequently, these cells were released from thymidine block and then allowed to undergo mitosis. The newly synthesized SNAP-Nup188 was labeled with TMR-star dye (yellow) at each timepoint shown. *α*-Pericentrin labeling the centrosome is shown in magenta. DNA was visualized by Hoechst staining (blue). n.a. is not applicable. Scale bar is 5 μm. Magnifications of boxed regions encompassing centrosomes in red, yellow and magenta is shown on right. Scale bar is 0.5 μm. Bottom panels show background fluorescence in cells not expressing SNAP-Nup188 (-dox).

Knowing that SNAP-Nup188 mirrors the behavior of Nup188, we pulse-labeled S-phase synchronized cells with a SNAP-binding 647-SiR dye. At timepoints after release from S-phase, we then chase-labeled SNAP-Nup188 with a TMR-Star dye. Thus, we followed two pools of SNAP-Nup188: an existing “old” pool and one that is derived from new synthesis. Interestingly, by monitoring total levels of SNAP-Nup188 (old) and SNAP-Nup188 (new) we observed that the old pool was unchanged as cells progressed through mitosis (Figure 2B, C). In contrast, the newly synthesized pool appeared to peak and then decline after mitosis. Therefore, only SNAP-Nup188 that is newly synthesized is subject to mitotic oscillation (Figure 2B, C).

That there might be two distinct pools of Nup188 correlates with the observation that Nup188 exists at NPCs and at centrosomes. To understand whether the centrosomal Nup188 pool is populated by new synthesis or exchange from intact NPCs (during interphase) or NPC breakdown (during mitosis), we directly examined the localization of “old” and “new” SNAP-Nup188 in synchronized cells. Consistent with the steady-state distribution of Nup188, pulse-labeled SNAP-Nup188 (old) can be found at both NPCs and at centrosomes (see *α*-Pericentrin labeling; Figure 2D, 0 h panel). Interestingly, by 4 h post-release from the S-phase-block, we no longer detected SNAP-Nup188 (old) at centrosomes whereas the pool at NPCs remained unchanged. These data suggest that centrosomal Nup188 is not populated by exchange from NPCs during interphase and that it likely turns over. In contrast, after just one hour of release from S-phase arrest, we observed a striking accumulation of SNAP-Nup188 (new) that was specific to centrosomes (Figure 2D). This was surprising because most nups that associate with the mitotic apparatus do so after release from the NPC during nuclear envelope breakdown (Belgareh et al., 2001; Cross and Powers, 2011; Joseph et al., 2004; Loiodice et al., 2004; Lussi et al., 2010; Wong et al., 2006). By 4 h a nuclear rim was also visible, but note that there are high levels of background fluorescence (see -dox panel, bottom) under these TMR-Star dye-labeling conditions that make it less obvious than expected. Nonetheless, these data support that newly synthesized Nup188 is targeted to centrosomes where it might then be turned over. Moreover, as cells exited mitosis, we could only detect Nup188 (new) at centrosomes, despite the fact that Nup188 (old) would have had access to the centrosome during mitotic NPC breakdown (Figure 2D, 12 and 16 h panels). These data suggest that once licensed to NPCs, Nup188 cannot bind to centrosomes, which can only be populated by a virgin pool.

### Nup188 distribution at steady-state is controlled by differential turnover

That new synthesis and turnover could control Nup188 localization at centrosomes prompted us to examine both of these processes in isolation. To examine transcription, we performed quantitative (q) RT-PCR on total mRNA generated from samples arrested in S-phase and released into mitosis (Figure 3A). As expected, we observed ∼4 fold higher *CYCLIN B* mRNA levels when M-phase (12 h after S-phase release) and S-phase samples were directly related (Figure 3A). We performed a similar RT-qPCR analysis of additional nup messages. Surprisingly, as shown in Figure 3A, we observed little change in nup transcripts with only *NUP35/53* mRNA and *NUP62* mRNA showing ∼2-fold increases in mitotic extracts. *NUP188* mRNA, by contrast, was unchanged. Thus, it is likely that the observed increase in Nup188 levels during mitosis is due to an inhibition of its degradation, not transcriptional regulation. This idea would also explain the observation that the SNAP-Nup188 protein is similarly regulated despite being controlled by another promoter (Figure 2A).

**Figure 3.**
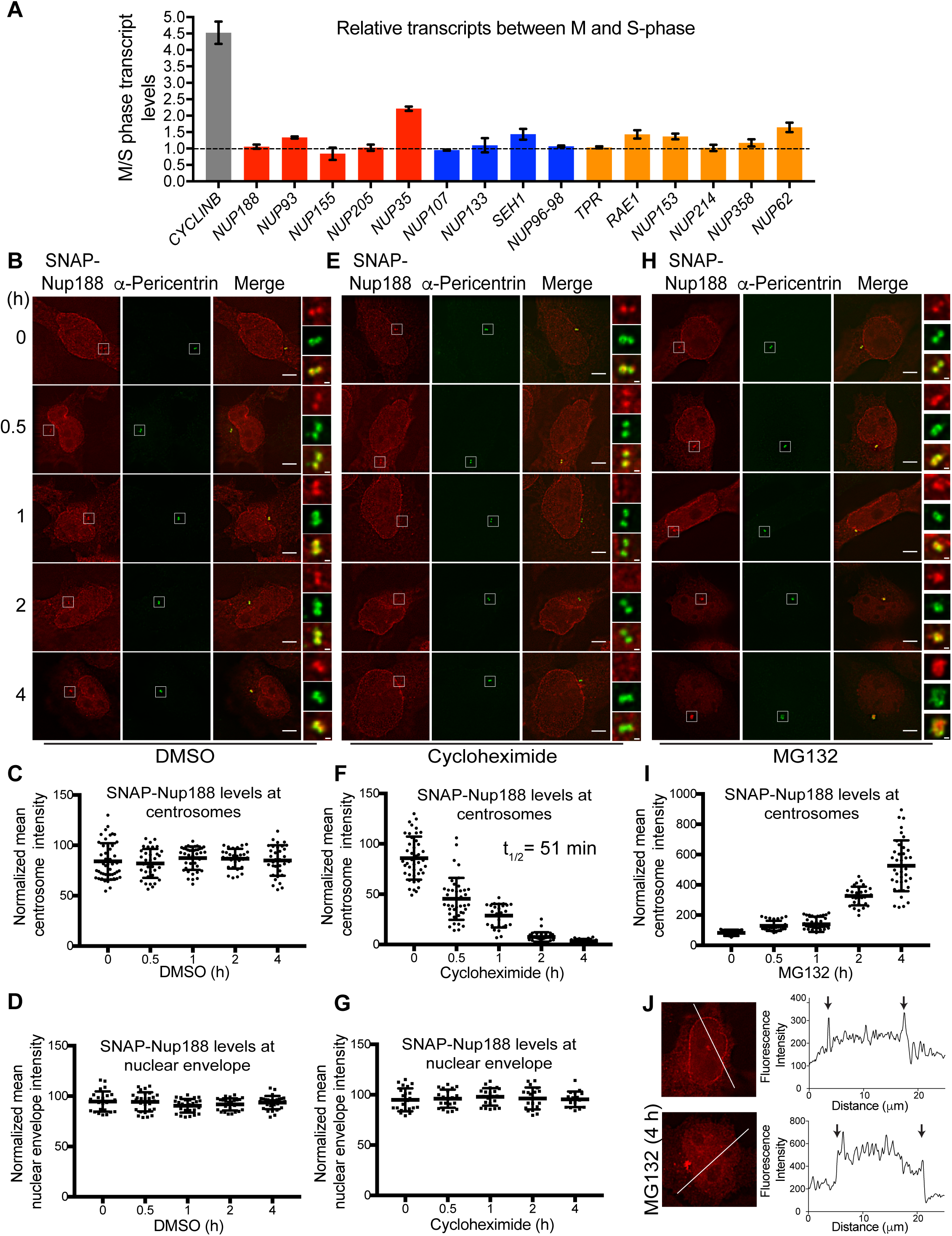
Nup188 turns over rapidly at centrosomes but is stable at NPCs. **(A)** *NUP188* mRNA levels are unchanged between S- and M-phase. Plot shows transcript levels at M-phase relative to S-phase levels as determined by RT-qPCR of the indicated nup messages. Shown are mean transcript levels (+/− SD) from three independent experiments. Dotted line reflects a ratio of 1, which indicates no change in relative levels between these cell cycle stages. Red, blue and orange are inner ring, outer ring and a mixture of cytoplasmic filament/nuclear basket nup genes, respectively. **(B)** Asynchronous HeLa cells producing SNAP-Nup188 were treated with DMSO (carrier-only control) and then labeled with 647-SiR dye (red) at indicated time points after drug addition, before imaging by fluorescence microscopy. Centrosomes were labeled with *α*-Pericentrin (green). Bar is 5 μm. Magnifications of boxed regions encompassing centrosomes in red, green and merge shown on right. Scale bar is 0.5 μm. **(C)** A representative plot (one of three independent replicates) of the mean intensity (+/− SD) of SNAP-Nup188 fluorescence at centrosomes in individual cells from experiment in (B) over time. **(D)** As in C but plotting nuclear envelope fluorescence. **(E-G)** As in A-D but cells were treated with cycloheximide. **(H-I)** As in A-C but cells were treated with MG132. **(J)**. Example fluorescence images of cells expressing SNAP-Nup188 treated with MG132 for 0 or 4 h. Line profiles of the fluorescence along white lines shown in images bisecting the nucleus shown in right panels. Note the y-axis scale and intranuclear fluorescence in bottom panels. Arrows denote location of nuclear rim along line profile plot.

Consistent with the idea that Nup188 levels at centrosomes are controlled by degradation, we treated cells with the proteasome inhibitory MG132 and monitored Nup188 by Western blot in cells synchronized and released from mitosis. As shown in Figure S1B, MG132-treatment led to a stabilization of Nup188 for several hours after release from M-phase-arrest when compared to carrier-alone (DMSO) samples, which declined as expected. Further, we could detect an increase of K48-linked ubiquitin-species associated with immunoprecipitations of FLAG-Nup188 specifically in MG132-treated cells (Figure S1C). Thus, it is likely that at least a portion of Nup188 is degraded at the end of mitosis. However, as Cyclin B is also stabilized under these conditions, we could not fully rule out that MG132-mediated mitotic arrest was simply upstream of Nup188 turnover.

We therefore more directly examined SNAP-Nup188 turnover at both NPCs and centrosomes outside of any mitotic regulation. Asynchronous cells were treated with cycloheximide to inhibit all protein synthesis and the levels of SNAP-Nup188 fluorescence were monitored over 4 h by labeling with a 647-SiR dye at each time point. As a control, we first measured the fluorescence of SNAP-Nup188 at the nuclear envelope and at centrosomes in carrier (DMSO) treated cells, which remained unchanged in both locations over this time period (Figure 3B-D). In cycloheximide-treated cells, fluorescence of SNAP-Nup188 at the nuclear envelope also remained constant over 4 h, which is consistent with the idea that Nup188 does not appreciably turnover once integrated into NPCs (Figure 3E, G). In contrast, we observed the complete loss of SNAP-Nup188 fluorescence at centrosomes between one and two hours (t_1/2_ of 51 min, Figure 3E, F). Further, to confirm that this loss was due to proteasome-mediated degradation, we also treated cells with MG132 (Figure 3H). Here, we observed a striking increase in SNAP-Nup188 fluorescence at centrosomes to levels ∼5 times higher than those of DMSO-treated samples (note change in y-axis scale, Figure 3I), alongside a concomitant increase in *α*-Pericentrin staining. Interestingly, we also observed an increase in SNAP-Nup188 fluorescence within the nucleus, which overshadowed and precluded our ability to directly assess specific nuclear envelope fluorescence under these conditions (compare line profiles of 0 and 4 h images; Figure 3J). Nonetheless, we interpret these data in a model in which proteasomal degradation of Nup188 is the major determinant of Nup188 levels at centrosomes.

### Nup188 is found at the inner layer of the PCM

To further substantiate that Nup188 is a bona fide component of centrosomes, even during interphase, we turned to super-resolution microscopy, which has revealed that the interphase centrosome is organized in sequential concentric layers that extend radially out to ∼200 nm from the centriole core (Fry et al., 2017; Lawo et al., 2012; Mennella et al., 2012; Sonnen et al., 2012). We labeled SNAP-Nup188 with a TMR-Star dye and performed structured illumination microscopy (3D-SIM) while co-labeling the inner (*α*-Cep192), intermediate (*α*-Cep152) and outer (*α*-Pericentrin) centrosome layers (Figure 4D) (Lawo et al., 2012; Mennella et al., 2012; Sonnen et al., 2012). As shown in Figure 4A, SNAP-Nup188 fluorescence appeared in a circular-pattern typical of PCM components when visualized down the long axis (top-view; Figure 4D) of the centriole. All three established PCM labels exhibited a qualitatively similar distribution, although unlike Cep152, which is only found at the proximal ends of centrioles (Cizmecioglu et al., 2010; Lukinavicius et al., 2013; Sir et al., 2011; Sonnen et al., 2013), SNAP-Nup188 coated the entire centriole length (see side-view, Figure 4B).

**Figure 4.**
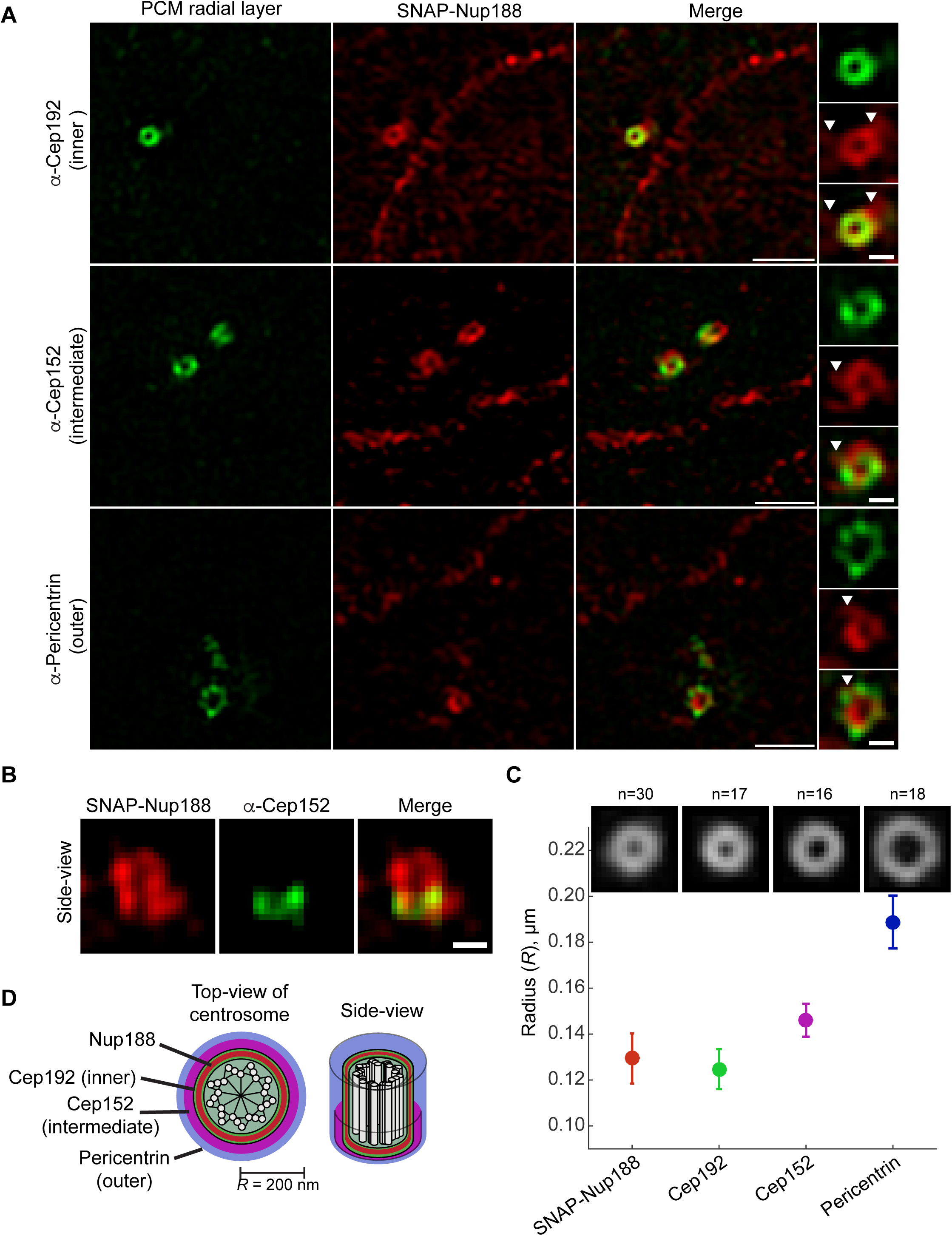
Nup188 is a component of the inner PCM layer. **(A)** Micrographs generated using 3D-SIM showing the localization of SNAP-Nup188 (labeled with TMR-Star; red) co-labeled (in green) with: *α*-Cep192, *α*-Cep152 and *α*-Pericentrin marking inner, intermediate and outer radial layers of the interphase PCM, respectively. Scale bar is 1 μm. Boxed regions encompassing the centrosomes showing green, red and merge panels are magnified at right. Scale bar is 0.25 μm. **(B)** 3D-SIM image of a side-view of SNAP-Nup188 and *α*-Cep152 labeling of centrosome. Scale bar is 0.25 μm. **(C)** Plot showing the quantification of the radius (*R*) of the centrosomal ring for the indicated proteins based on a ring structure fitting to individual images (*n* indicates number of images from three independent replicates). Mean (solid circle) and SD (bars) are shown. **(D)** Approximate-scale schematic of centrosome showing centriole in center (white circles are microtubules) and location of Nup188 within the inner layer.

To gain quantitative insight into SNAP-Nup188’s radial distribution, we fitted a theoretical circle to individual SNAP-Nup188 images and used these results to generate an average image (Figure 4C). This analysis showed that the average radius of SNAP-Nup188 fluorescence was 129 nm, which placed it closest to the inner layer demarked by Cep192 (*R* of 125 nm)(Figure 4C). Consistent with this, SNAP-Nup188 was clearly found inside of both Cep152 (*R* of 146 nm) and Pericentrin (*R* of 189 nm)(Figure 4C). Thus, Nup188 is found within the inner layer of the interphase centrosome, closest to the centriole. This observation predicts that Nup188 might physically interact with inner-layer components like Cep192.

Interestingly, unlike Cep192 and Cep152, SNAP-Nup188 was also found in extensions that emanate outward from its circular core that often intercalated into Pericentrin (Figure 4A, arrowheads). These extensions are particularly obvious when comparing montages of SNAP-Nup188 images to those of either *α*-Cep192 or *α*-Cep152 (Figure S2A-D). We wondered whether these might reflect interactions with microtubules. We therefore de-polymerized microtubules using nocodazole but found little change to these structures (Figure S2E). Nonetheless, these data further support the conclusion that Nup188 is a component of PCM associated with the inner centrosome layer and likely other, yet to be defined, structures.

### Proximity-labeling uncovers PCM-specific interactors

Having more clearly established that Nup188 has the characteristics of a component of PCM, we sought to define the molecular interactions that contribute to this association. We therefore turned to a biotin-ligase proximity-labeling approach (Roux et al., 2012) to avoid the challenges with extracting Nup188 from two relatively insoluble protein assemblies (i.e. NPCs and centrosomes). Moreover, as Nup188 turns over rapidly at centrosomes, we wanted to exploit that this approach reflects a “history” of proximity associations, thus increasing the likelihood that we would find specific centrosome interactors. To reduce background biotinylation, we generated a stable HeLa cell line that expresses the promiscuous biotin ligase (BirA*)(Roux et al., 2012) both alone and fused to Nup188, expressed behind a dox-inducible promoter (Figure 5A). We then titrated dox concentrations to produce below-endogenous Nup188 levels (Figure S3A). To ensure the specificity of BirA*-Nup188 labeling, we stained cells with fluorescently labeled (Alexa Flour™ 488) streptavidin. Gratifyingly, we detected streptavidin-labeling at centrosomes (indicated by *α*-γ-Tubulin, red) in interphase and mitosis, and at the nuclear periphery (in interphase)(Figure 5B, upper panels, maximum intensity projections shown). We did not detect any streptavidin-labeling of either centrosomes or NPCs when BirA* was expressed on its own (Figure 5B, lower panels). Consistent with these observations, probing total proteins separated by SDS-PAGE and transferred onto nitrocellulose with streptavidin coupled to HRP showed a distinct subset of biotinylated proteins between BirA* and BirA*-Nup188 cells (Figure 5C).

**Figure 5.**
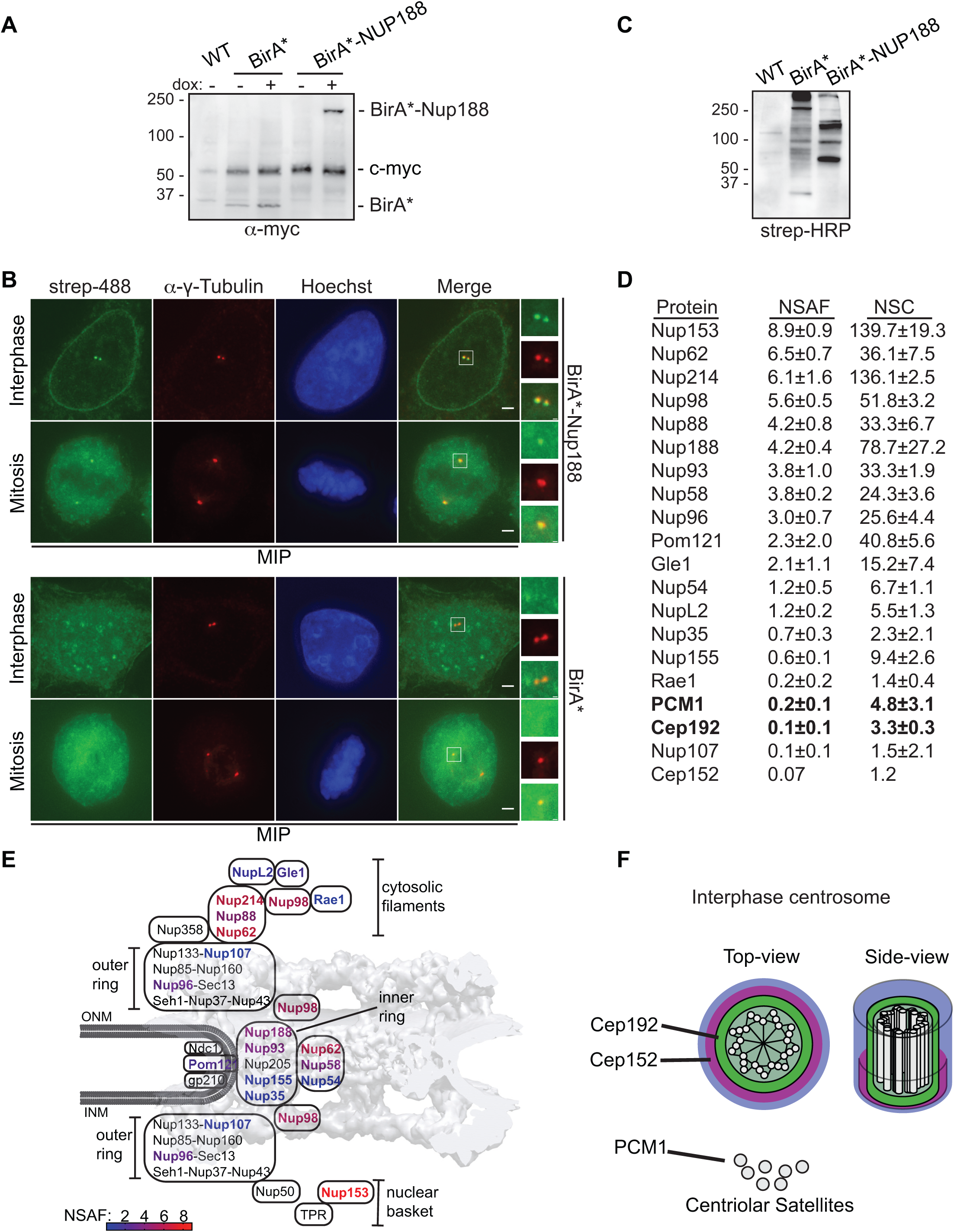
Proximity labeling determines physical interactions between Nup188 and PCM. **(A)** BirA* and BirA*-Nup188 are stably and inducibly-expressed at similar levels. Western blots of whole cell extracts from BirA* and BirA*-Nup188-expressing cell lines. Both BirA* and BirA*-Nup188 are under the control of a doxycycline (dox) - inducible promoter and produced with a myc epitope to allow detection by Western blot. Note that endogenous c-myc is also detected in all lanes and serves as a total protein load reference. Numbers at left are positions of molecular weight standards (in kD). **(B)** Centrosomes and the nuclear envelope are specifically biotinylated in cell lines expressing BirA*-Nup188. Maximum-intensity projections (MIP) of a z-series of fluorescence micrographs of BirA*-Nup188 (top panels) and BirA* alone (bottom panels) expressing cells. DNA is stained with Hoechst to help identify metaphase cells and visualize the nucleus. Biotinylated proteins are labeled with streptavidin (strep) coupled to Alexa-488 (green). Centrosomes are labeled with *α*-γ-Tubulin (red). Scale bar is 5 μm. Boxed regions of centrosomes in green, red and merge are magnified at right. Scale bar is 1 μm. **(C)** Total protein from HeLa cells (WT) and those producing BirA*and BirA*-Nup188 was separated by SDS-PAGE; biotinylated proteins detected by streptavidin (strep) coupled to HRP. Numbers at left are positions of molecular weight standards (in kD). **(D)** Table of biotinylated proteins identified in BirA*-Nup188-expressing cells with NSAF and normalized spectral count (NSC) values (mean ± SD) from three independent replicates. See Table S1 for all proteins identified. **(E)** Schematic of the relative location of nups within a diagram of the NPC superimposed on a cryo-EM map of the NPC structure (von Appen et al., 2015). Biotinylated nups are color-coded based on the heat-map scale of NSAF values indicated at bottom. **(F)** Approximate scale diagram of centrosome showing position of Cep192 and also PCM1 as a component of centriolar satellites.

Confident in the specificity of the biotinylation in our cell lines, we affinity purified biotinylated proteins using streptavidin magnetic beads from both BirA*-Nup188 and BirA*-alone expressing cell extracts. Protein eluates were subjected to trypsin digestion followed by LC MS/MS peptide identification (Table S1). As the majority of Nup188 is associated with NPCs, most identified peptides corresponded to nups. We normalized specific peptide abundance to the total peptides identified in a given experiment and to the length (in amino acids) of the protein, which we present as a Normalized Spectral Abundance Factor (NSAF) value (Figure 5D). We have also color-coded nup names in a heatmap, which we superimpose on the NPC structure to give a sense of the location of the biotinylation hotspots in the NPC (Figure 5E). As expected, based on Nup188’s established location within the inner ring complex, we detected Nup93 as one of the top interactors. Consistent with this, Nup62 and Nup58 were also identified suggesting that the BirA* enzyme is likely proximal to the N-terminus of Nup93 where these FG-nups connect to the inner ring (Stuwe et al., 2015). Interestingly, Nup98 has a high NSAF value consistent with other data supporting that Nup188 binds to this FG-nup (Fischer et al., 2015; Onischenko et al., 2017). Further, the highest NSAF values were found to be for Nup153 and Nup214, components of the nuclear basket and cytoplasmic filaments, respectively. These data suggest that Nup188 might function in several locations in the NPC and/or associate with these nups in other subcellular locations.

We classified the other ∼80 proteins identified (see complete lists, Table S1), using Gene Ontology-function analysis (Figure S3B), which suggested interactions with many nuclear components including RNPs and chromatin-binding proteins. Within this sea of proteins, only two are components of PCM, Cep152 and Cep192, and one (PCM1) is a component of both centriolar satellites and PCM (Figure 5D, F, Table S1). Of these, Cep192 and PCM1 were found in all three biological replicates with a mean NSAF value of 0.1 and 0.2, respectively. While this is expectedly low (due to the overall low abundance and rapid turnover of Nup188 at centrosomes versus NPCs), they were remarkably specific as no other centrosome components were detected. Thus, these data further support that Nup188 is found in the inner layer of the PCM in proximity to Cep192. To help explain the interaction with PCM1, we tested whether SNAP-Nup188 came into proximity to PCM1 at centriolar satellites or centrosomes. As shown in Figure S2E, we observed a few instances where PCM1 foci at centrosomes were adjacent to or co-localized with SNAP-Nup188 by 3D-SIM in a microtubule-independent fashion. These data at least provide a rationale for why PCM1 was detected by BioID.

### Specific knockdown of Nup188 impacts centriolar satellite distribution

That Nup188 interacts with Cep192 and PCM1 raised the possibility that these proteins might be required for its recruitment to centrosomes. We therefore used siRNA to knockdown Cep192, Cep152 and PCM1 and examined their impact on SNAP-Nup188 localization (Figure S4A). Importantly, all three of these proteins were depleted with the siRNA treatments (Figure S4B). However, despite clear reductions in the specific proteins, we were unable to convincingly observe any changes to SNAP-Nup188 localization (Figure S4A) with mean values of SNAP-Nup188 fluorescence remaining unchanged (Figure S4C). These data suggest that Nup188 might make multiple connections with the centrosome inner layer beyond just Cep192. Likewise, depletion of Nup188 did not impact the recruitment of either Cep192 or Cep152 to centrosomes (Figure S5A, B), despite the fact that we could clear SNAP-Nup188 from centrosomes with siRNA treatments as short as 12 h due to its rapid turnover rate (Figure S6A-C).

While knockdown of Cep192 and PCM1 was insufficient to upset Nup188 PCM accumulation, we nonetheless assessed whether Nup188 might impact centrosome function guided by the established roles of these two proteins. First, as PCM1 is a major component of centriolar satellites, we tested the distribution of PCM1 in the context of siRNA knockdown of *NUP188*. Strikingly, PCM1-puncta were no longer distributed in a close-radial association with centrosomes but instead were dispersed throughout the cytosol (Figure 6A). To quantitatively assess PCM1 distribution around centrosomes, we calculated the total fluorescence of *α*-PCM1 staining as a function of its radial distribution from the centrosome axis (Figure 6B). Cells treated with *NUP188* siRNA had, on average ∼30% less fluorescence compared to control cells treated with scrambled siRNA at radii within 5 μm of the centrosome (Figure 6A, B; Figure S7A). Importantly, the reduction of *α*-PCM1 fluorescence reflected the dispersal of PCM1, not its disappearance, as shown in Figure 6C (and Figure S7B) where we have plotted the cumulative distribution in the same cell areas: PCM1 foci are more likely to populate cytosol further from the centrosome axis in cells treated specifically with *NUP188* siRNA. Concerned that this effect on satellites might be due to a disruption of nuclear transport, we also tested knockdown of several additional nups including Nup93, Nup62 and Nup214. Over the time course of this experiment, we observed similar levels of knockdown for each of these proteins (Figure S7C) and yet, only the knockdown of Nup188 had any significant impact on PCM1 distribution. Indeed, even reduction of Cep192 levels did not influence PCM1 localization (Figure 6B, C and Figure S7A, B). Thus, when considered in the context of evidence for a physical interaction, we suggest that this effect may be a direct consequence of perturbing a functional relationship between Nup188 and PCM1.

**Figure 6.**
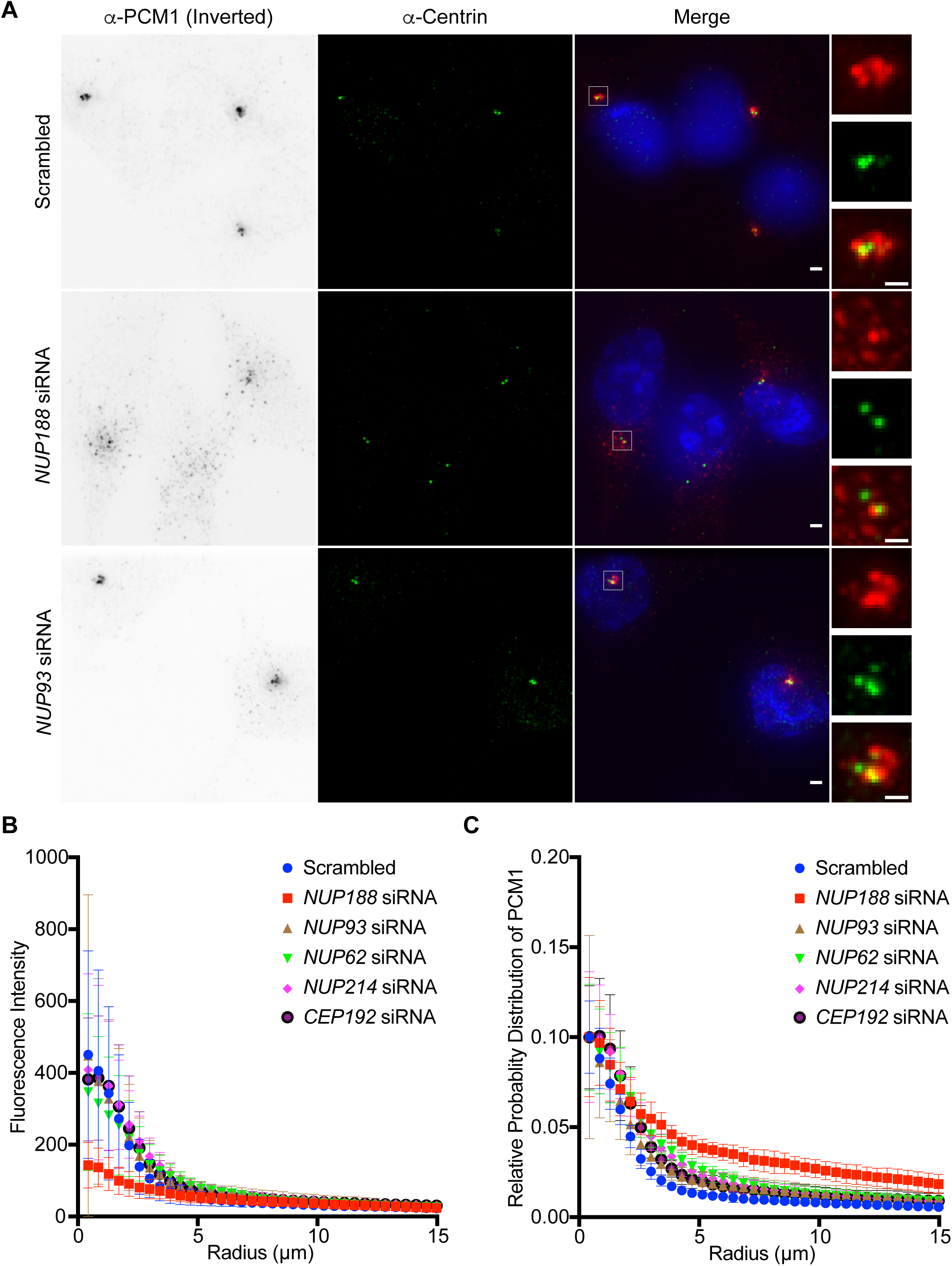
Knockdown of *NUP188* specifically impacts centriolar satellite distribution. **(A)** Fluorescence images of U2OS cells labeled with α-PCM1 (fluorescence inverted) and *α*-Centrin (green) to localize centrioles. Cells were treated for 48 h with siRNAs specific to indicated nup transcripts with scrambled siRNA control before imaging. Scale bar is 5 μm. Boxed regions of centrosomes in green, red and merge are magnified at right. Scale bar is 2.5 μm. **(B)** Plot of the total fluorescence of *α*-PCM1 labeled centriolar satellites within an annulus of indicated radii around an axis of *α*-Centrin fluorescence in 10 cells each treated with the indicated siRNA. Error bars indicate ±SD. Another experimental replicate can be found in Figure S7A. **(C)** As in (B) but plotting the cumulative distribution of *α*-PCM1 labeling throughout the cell (n=10). Error bars indicate mean ±SD. Another experimental replicate can be found at Figure S7B.

### Nup188 is required for centriole duplication through Sas6 loading

We next explored how Nup188 might impact Cep192 function, which has been ascribed to diverse roles in the centrosome lifecycle including in centriole duplication (Kim et al., 2013; Sonnen et al., 2013; Zhu et al., 2008) and centrosome maturation (Gomez-Ferreria et al., 2007; Joukov et al., 2014; O’Rourke et al., 2014; Zhu et al., 2008). We therefore began by testing the impact of knockdown of Nup188 or other nups including Nup93, Nup62 and Nup214, on centriole duplication. Centriole number was assessed by immunostaining with either *α*-Centrin or *α*-CP110 antibodies, which recognize the distal ends of centrioles. Here, the results were again remarkably specific for Nup188 (Figure 7A, B). For example, knockdown of Nup62 and Nup93 did not lead to statistically different changes in centriole number with ∼50% of cells having between 2-4 centrioles like the scrambled siRNA-treated control. Small fractions of each of these cell populations also had supernumerary centrioles, which were particularly striking in Nup214-depleted cells (Figure 7A, B). In contrast, only 20% of *NUP188* siRNA-treated cells had between 2-4 centrioles with the other 80% containing 0 to 2. Thus, knockdown of Nup188 leads to a specific abrogation of centriole duplication.

**Figure 7.**
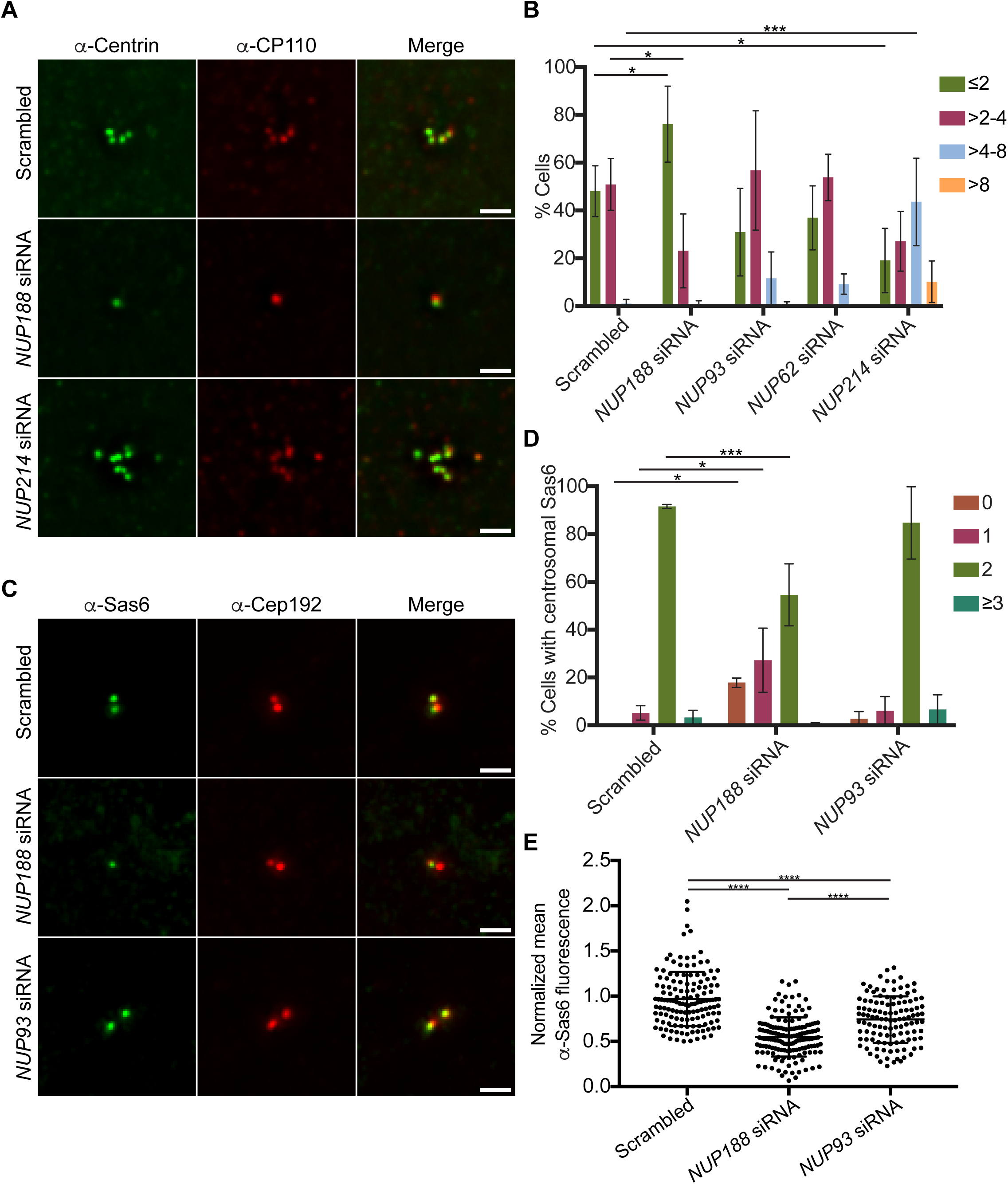
Nup188 is required for centriole duplication through Sas6 loading. **(A)** Fluorescence micrographs of U2OS cells treated with the indicated siRNAs. Centrioles are labeled with both *α*-Centrin (green) and *α*-CP110 (red). Scale bar is 5 μm. **(B)** Plot of the quantification of centriole number (*α*-Centrin foci) in U2OS cells treated with the indicated siRNAs. Three independent replicates of >30 cells per replicate were quantified. Error bars indicate mean ±SD. *P*-values are calculated from two-way ANOVA with Dunnett’s multiple comparison test. **(C)** Fluorescence micrographs of S-phase-synchronized U2OS cells stained with *α*-Sas6 (green) and *α*-Cep192 (red) antibodies. Scale bar is 5 μm. **(D)** Plot of the percentage of cells with the indicated number of Sas6 foci in S-phase synchronized U2OS cells treated with the indicated siRNAs. Three independent replicates of >40 cells per replicate were quantified. Error bars indicate mean ±SD. *P*-values are calculated from two-way ANOVA with Dunnett’s multiple comparison test. **(E)** Plot of mean fluorescence (normalized to intracellular background) of individual *α*-Sas6 foci. Three independent replicates of >35 cells per replicate were quantified. Error bars indicate mean ±SD. *P*-values are calculated from one-way ANOVA with Tukey’s multiple comparison test.

Finally, to gain further insight into the mechanism of the Nup188-knockdown-specific centriole loss, we tested whether Nup188 helped control the recruitment of Sas6, which is thought to be critical for formation of the cartwheel. Because Sas6 is recruited during a short temporal window in S-phase (Leidel et al., 2005; Strnad et al., 2007), we arrested cells in S-phase using aphidicolin, which was confirmed by FACS analysis of DNA content (Figure S8). Under these conditions, reducing levels of Nup188 but not Nup93, led to a marked loss of Sas6 recruitment to centrosomes (*α*-Cep192, red) with ∼50% of cells showing aberrant number (0 and 1) of Sas6 foci on centrosomes, thus providing a rationale for the centriole duplication defect (Figure 7C, D). Indeed, even in cases where we observed Sas6 at centrosomes, the *α*-Sas6 labeling fluorescence was clearly reduced suggesting that fewer Sas6 molecules are recruited in the absence of Nup188 (Figure 7E). Thus, Nup188 is specifically required for centriole duplication, likely through a role contributing to Sas6 recruitment and/or stability.

## Discussion

Here, we argue that we have made a definitive case for Nup188 being a component of the centrosome. There are several reasons for this assertion: 1) Endogenous Nup188, but also SNAP- and BirA*-fusions of Nup188 reproducibly and unambiguously localize to centrosomes; this localization is not dependent on microtubules as it is stable under conditions of microtubule depolymerization. 2) We establish physical links between Nup188 and centrosome components, namely Cep192 and PCM1. 3) We place Nup188 within the PCM in a physical context that is consistent with it interacting with Cep192 (i.e. within the centrosome inner layer) 4) Like other PCM components, Nup188 is further enriched at centrosomes during its maturation in mitosis. 5) We demonstrate the specific disruption of centriole duplication upon Nup188, but not other nup, knockdown.

One of the more interesting aspects of Nup188’s association with centrosomes is that it is recruited at all stages of the cell cycle. For example, other nups like the Nup107-160 complex are recruited to kinetochores solely during mitosis, presumably from a pool liberated during NPC breakdown (Belgareh et al., 2001; Loiodice et al., 2004). In contrast, mechanisms must be in place to sort newly synthesized Nup188 to at least two different destinations throughout interphase. Once there, the levels of Nup188 accumulation are ultimately controlled by differential turnover with the NPC pool being extremely stable and the centrosome pool sensitive to proteasomal degradation. It will be interesting to explore the functional implications of these observations. For example, one prediction would be that the centrosomal pool would be much more sensitive to siRNA knockdown (which it is, Figure S6) and perhaps to overexpression. Thus, this might help to explain why disruption of the PCM function of Nup188 is more sensitive to partial knockdown in our heterotaxy *Xenopus* model over its function at NPCs (Del Viso et al., 2016). It might also predict that putative Nup188 disease variants that destabilize Nup188 would be more likely to affect its PCM function as well.

Thus, it seems reasonable to conclude that the modulation of the Nup188 turnover rate is the major determinant of its accumulation at centrosomes during mitosis. The expectation is that Nup188’s turnover would be inhibited during G2 and early M-phase to elevate its levels, and then stimulated at the end of mitosis to trigger its degradation. While we have not yet identified the molecular players that control this behavior, clear candidates are E3 ubiquitin ligases like the anaphase-promoting complex (APC) or the SKP-CUL1-F Box (SCF) complex, which have both been implicated in centrosome function by impacting the degradation of PCM components (Arquint et al., 2018; Cunha-Ferreira et al., 2009; D’Angiolella et al., 2010; Drosopoulos et al., 2014; Freed et al., 1999; Holland et al., 2010; Piva et al., 2002; Prosser et al., 2012). Other E3 ligases have also been implicated to function at centrosomes including Mib1 (Cajanek et al., 2015). Regardless, these data all point to proteasomal turnover as a major determinant of centrosome architecture and size. This idea is best illustrated by the observation that inhibiting proteasomes during interphase can lead to over a 5-fold increase in the levels of Nup188 at centrosomes. Indeed, this appears to be a general rule for at least a subset of PCM components (Wigley et al., 1999) and reinforces that Nup188 is a component of PCM.

Further illustrating the central importance of controlling the degradation of centrosomal proteins in time and space, even Plk4 and Sas6 levels at centrosomes, and thus the centriole duplication mechanism, is under the control of the ubiquitin-proteasome system (Arquint and Nigg, 2016; Cunha-Ferreira et al., 2009; Holland et al., 2010; Puklowski et al., 2011). A priority going forward therefore is to fully elucidate how Nup188 functions during centriole duplication and whether there is a role for Nup188 turnover in this process as well. Our data support a model in which Nup188 impacts events at or upstream of Sas6 recruitment to give rise to the procentriole. While we did not identify Sas6 by BioID, this is not surprising as it is only recruited to centrioles for a short time period during G1-S before being degraded (Leidel et al., 2005; Strnad et al., 2007). Nonetheless, even if we cannot be conclusive as to whether Nup188 biochemically interacts with Sas6, we favor models in which it acts through a mechanism analogous to Cep192, which provides a platform for Plk4 responsible for Sas6 recruitment (Arquint and Nigg, 2016; Kim et al., 2013; Sonnen et al., 2013). Certainly, as with many nups, Nup188 is a target of kinases with several predicted potential polo kinase sites (Gouw et al., 2018), suggesting this as a likely possibility for future investigation.

Centrosomal Nup188 likely makes several direct connections with proteins beyond PCM1 and Cep192, as depletion of either did not impact Nup188 recruitment to this organelle. These results illustrate the persistent challenge of delimiting a function for Nup188 at centrosomes in a context in which the perturbation of NPC function can be fully ruled out. Here, we nonetheless make the case that Nup188 at centrosomes directly contributes to mechanisms that control both centriolar satellite distribution and centriole duplication. In addition to the physical and functional links that we have uncovered between Nup188 and these processes, this assertion is further supported by the remarkable specificity of Nup188 knockdown in mediating these centrosomal-related defects over even its nearest neighbor binding partner (at the NPC), Nup93. These results predict that Nup93 is not a component of centrosomes and provides one of a handful of examples where one nup might separate from its nup subcomplex to perform a unique function. Another example being Nup96, which interestingly is also specifically degraded during mitosis while the rest of the outer ring complex remains stable (Chakraborty et al., 2008). A model in which Nup188 is disengaged from its binding partner Nup93 at centrosomes could also help to explain why the NPC-assembled pool of Nup188 (which would be bound to Nup93) does not seem able to populate centrosomes even after “solubilization” by NPC breakdown during mitosis. Such a model needs to be reconciled, however, with our prior data suggesting that both Nup93 and Nup188 can localize to cilia bases (Del Viso et al., 2016).

Further work still remains to define the catalogue of nups at centrosomes (and at cilia bases)(Breslow et al., 2013; Del Viso et al., 2016; Endicott et al., 2015; Kee et al., 2012) with key candidates being the FG-nups Nup62 and Nup98, both of which show analogous, if more striking, cyclin-like behavior during mitosis. Further, each has been implicated in some form of mitotic regulation: Nup62 has a potential pool at centrosomes (Hashizume et al., 2013) and Nup98 potentially directly impacts spindle assembly (Cross and Powers, 2011) while also localizing to cilia bases (Endicott et al., 2015; Endicott and Brueckner, 2018). Could Nup188 interface with these FG-nups in the PCM? Recent work, for example, supports that Nup188 and its paralog Nup205 have atomic structures that resemble the karyopherin/importin β-family of NTRs (Andersen et al., 2013; Flemming et al., 2012; Sampathkumar et al., 2013; Stuwe et al., 2014). In fact, evidence indicates that these interactions might be critical for helping to hold the NPC structure together (Onischenko et al., 2017) perhaps by “ordering” the inherently disordered FG-repeats, which themselves have been shown to display a spectrum of collective behaviors and “phases” (Lemke, 2016) some of which are impacted by NTR-binding (Hofweber et al., 2018; Yoshizawa et al., 2018). Could an NTR-like Nup188 also contribute to an ordering or disordering of centrosome organization? Such a hypothesis becomes compelling as the *C. elegans* SPD-5 is capable of undergoing a phase-change to a gel/condensate-like state required for microtubule polymerization (Woodruff et al., 2017). That the more structured layers of the interphase centrosome are lost as the centrosome matures with the addition of more PCM (including Nup188) before mitosis (Haren et al., 2009; Khodjakov and Rieder, 1999; Lawo et al., 2012; Lee and Rhee, 2011; Sonnen et al., 2012; Zhu et al., 2008), suggests that similar changes might occur to the human centrosome as well. We look forward to directly examining whether Nup188 impacts this process in future work.

## Materials and Methods

### Plasmids

Plasmids used in the study are listed in Table S5. To construct pMC1 and pMC2, the GFP ORF from GFP-NUP188 plasmid (Gift from Dr. K. Tanaka (Itoh et al., 2013)) was replaced with either *SNAP* or *BIRA\** coding sequences excised from pSNAPf (NEB) and pcDNA3.1 mycBioID plasmid (Addgene), respectively. To generate pNV2 and pNV3, SNAP-Nup188 and BirA*-Nup188 were sub-cloned into pcDNA5/FRT/TO. To construct pNV1, *NUP188* sequence was excised from pNV2 plasmid leaving only the *BIRA\** ORF. To construct pKD1 the *NUP188* coding sequence was amplified by PCR and subcloned into p3XFLAG-CMV™-10 vector (Millipore Sigma) using the Gibson Assembly® Master Mix (NEB) according to the manufacturer’s instructions.

### Cell Culture, transfection and stable cell line generation

The following human cell lines were used in this study: HeLa-M (gift from Pietro De Camilli), HeLa (ATCC), HeLa T-Rex-Flp-In (Thermo Fisher Scientific) and U2OS (ATCC). HeLa, HeLa-M and HeLa T-Rex-Flp-In were cultured in Dulbecco’s Modified Eagle Medium (DMEM) with 4.5 g/L glucose (Thermo Fisher Scientific) and U2OS cells (ATCC) were cultured in McCoy’s 5A Medium Modified (Thermo Fisher Scientific). The medium was supplemented with 10% FBS (Millipore Sigma) and 100 units/ml penicillin and 100 μg/ml streptomycin (Thermo Fisher Scientific). The HeLa T-Rex-Flp-In cell line was also supplemented with 100 μg/ml Zeocin (Invivogen) and 15 μg/ml of Blasticidin (Invivogen) to maintain the stably integrated FRT site and tetracycline repressor, respectively. All cells were maintained in a humidified incubator at 37°C in 5% CO_2_.

Transfection of siRNA was carried out using RNAiMax (Thermo Fisher Scientific) as per the manufacturer’s instructions. Briefly, siRNAs were diluted in 500 μl of optiMEM (Thermo Fisher Scientific) to a final concentration of 30-50 nM (optimized by Western blot) in a 6 well plate. After mixing, 8 μl of RNAiMax was added and incubated for 20 min. Cells were trypsinized and diluted in 2 ml antibiotic-free medium and then added to the siRNA mix and incubated for 48 h, with the exception of Sas6 localization experiments, which were assessed after 72 h. Knockdown efficiency was assessed by Western blot. The siRNA duplex oligonucleotides are listed in Table S4.

To generate cell lines stably expressing BIRA*, BIRA*-NUP188 and SNAP-NUP188, a 9:1 molar ratio of pOG44 and pNV1, pNV2 and pNV3, respectively, were co-transfected into the HeLa T-Rex-Flp-In cell line. Two days after transfection, cells were selected for Hygromycin resistance by growing them in medium containing DMEM-high glucose (Thermo Fisher Scientific), 10% Tet system approved FBS (Takara), 100 units/ml penicillin and 100 μg/ml streptomycin (Thermo Fisher Scientific), 15 μg/ml of Blasticidin (Invivogen) and 200 μg/ml Hygromycin (Thermo Fisher Scientific). Culture medium was changed every three days. Clonal colonies were visible by 14 days, which were then isolated using 8 mm diameter cloning cylinders (Fisher Scientific), trypsinized, expanded and verified by Western blotting.

### Cell cycle synchronization and drug treatment

To establish S-phase synchronized cell populations, HeLa-M and SNAP-NUP188 cell lines were grown to 30% confluency and treated with 2 mM thymidine (Millipore Sigma) for 16 h. Cells were then washed twice with phosphate buffered saline (PBS) and incubated in medium without thymidine for 8 h. After this incubation, 2 mM thymidine (Millipore Sigma) was added to the medium for the next 16 h to block all cells in early S-phase. To enrich for cells in mitosis, S-phase-synchronized cells were released in the presence of 100 nM nocodazole (Millipore Sigma) for 12 h. Mitotic cells were then harvested by gently tapping the culture dish and collecting the medium. To investigate Nup188 turnover, HeLa cells expressing SNAP-NUP188 were treated with 10 μg/ml cycloheximide or 20 μg/ml MG132 or DMSO for up to 4 h at 37°C. To examine the recruitment of Sas6 to centrosomes, U2OS cells were arrested in S-phase by treatment with 2 μg/ml aphidicolin (Millipore Sigma) for 24 h. For microtubule-depolymerization experiments, cells were treated with 5 μg/ml nocodazole (Millipore Sigma) or DMSO for 3 h at 37°C.

### RNA Extraction, cDNA Synthesis and RT-qPCR

Comparison of expression levels of nup transcripts in synchronized cells was carried out by reverse transcription (RT) followed by quantitative (q) PCR. Briefly, cells were synchronized in S phase and released; cells were then harvested every hour. Total RNA was isolated using the RNeasy Plus mini kit (Qiagen) as per the manufacturer’s protocol. Superscript III RT kit (Thermo Fisher Scientific) was used to synthesize cDNA using Oligo-dT primers (Thermo Fisher Scientific). qPCR of the cDNA template was carried out by the iTaq Universal SYBR Green Supermix (Bio-Rad) with 0.2 pmol/μl of gene-specific primers (Integrated DNA Technologies)(Table S3). A region of the *GAPDH* cDNA was amplified as an internal control. The specificity of each amplicon was verified by a melt curve analysis.

### Western Blots

To extract total protein from whole cell extracts, cells were washed twice with PBS and then lysed in RIPA buffer (50 mM Tris, pH 7.4, 150 mM NaCl, 1% Triton X-100, 0.5% deoxycholate, 0.1% sodium dodecyl sulfate) supplemented with both protease and phosphatase inhibitor cocktails (Thermo Fisher Scientific) on ice for 10 min. Lysates were harvested using a cell scraper and then clarified by centrifugation for 10 min at 14,000 rpm at 4°C and supernatants were collected. Protein concentrations were determined by a Bradford assay (Thermo Fisher Scientific). 15-50 μg of protein was separated on a 4-20% gradient gel (Bio-Rad) and transferred onto 0.2 µm nitrocellulose (Bio-Rad) using the Mini Trans-Blot Cell (Bio-Rad) at 100 V for 60 min (Bio-Rad). Blocking and antibody incubations were performed with 5% milk (American Bio) in Tris-buffered saline, 0.1% Tween 20 (TBST). Nitrocellulose was blocked with 5% milk in TBST for 1 h, and then incubated with primary antibodies (listed in Table S2) for 1 h. Blots were then extensively washed in TBST and then incubated with horseradish peroxidase (HRP)-conjugated secondary antibodies for 1 h. After washing, proteins were detected using SuperSignal West Femto kit (Thermo Fisher scientific) and images were acquired with a ChemiDoc XRS+ Imaging (Bio-Rad) according to the manufacturer’s instructions.

### Immunofluorescence microscopy

To determine the localization of proteins in fixed cells by immunofluorescence microscopy, cells were first grown on coverslips, fixed in either −20°C methanol for 5 min or 4% paraformaldehyde (PFA) for 10 min at RT and washed twice with PBS. Cells were then blocked with either 5% FBS or 1% BSA in PBS with 0.1% Triton-X 100 (PBST). Cells were then incubated in primary antibody diluted in 1% FBS or 1% BSA in PBST for 1 h. After incubation cells were washed with PBS and then incubated with secondary antibody diluted in 1% FBS or 1% BSA in PBST for 1 h. To image biotinylated proteins, streptavidin-coupled to Alexa Fluor™ 488 (Thermo Fisher Scientific) was used. Lastly, cells were incubated with Hoechst (Thermo Fisher Scientific) to stain the DNA and coverslips were then mounted using Fluoromount-G™ (Electron Microscopy Sciences) before imaging.

Immunostained cells were imaged on a Deltavision widefield deconvolution microscope (Applied Precision/GE Healthcare) equipped with a 100X/1.40 (Olympus) or PLANAPO 60X/1.42 oil objective (Olympus), a CoolSnap HQ^2^ CCD camera (Photometrics) and Xenon arc lamp. Image stacks were acquired in 0.2 µm increments along the z-axis.

### SNAP-Nup188 labeling

To label SNAP-Nup188 with fluorophores, we used either a 647-SiR dye (SNAP-Cell® 647-SiR, NEB) or TMR-Star dye (SNAP-Cell® TMR-Star, NEB). Briefly, expression of *SNAP-NUP188* was induced by adding doxycyclin (Millipore Sigma) to the culture medium at a final concentration of 100 ng/ml for 48 h. Labeling was performed in living cells by addition of SNAP-Cell® 647-SiR or SNAP-Cell® TMR-Star directly to the culture medium at a final concentration of 2 μM. Labeling was performed for 30 min at 37°C. Cells were then washed three times in pre-warmed medium and further incubated in for 30 min at 37°C to remove excess unincorporated dye. After washing three times with PBS, cells were fixed and processed for immunofluorescence as described above.

To determine whether centrosomes were populated by newly synthesized Nup188 or from exchange from NPCs, SNAP-NUP188-expressing cells were synchronized in S-phase and pulse labeled with SNAP-Cell® 647-SiR. Subsequently, the cells were released from the S-phase block and chase-labeled every hour with SNAP-Cell® TMR-Star before processing for immunofluorescence microscopy.

To investigate the proteasomal-mediated turnover of SNAP-Nup188 at centrosomes, asynchronous cells were treated with either DMSO, cycloheximide or MG132 as described above for up to 4 h. Samples were labeled with SNAP-Cell® 647-SiR hourly intervals before being prepared for immunofluorescence.

### Structured illumination microscopy

To localize SNAP-Nup188 at centrosomes at subdiffraction resolution, we employed super-resolution three-dimensional structured illumination (3D-SIM). Cells producing SNAP-Nup188 were labeled with the TMR-Star dye, fixed with methanol at −20°C for 5 min and then immunostained with antibodies recognizing centrosome components. 3D-SIM imaging was performed on a DeltaVision OMX V3 system (GE Healthcare Life Sciences) equipped with a U-PLANAPO 60X/1.42 PSF oil immersion objective lens (Olympus, Center Valley, PA), CoolSNAP HQ2 CCD camera with a pixel size of 0.080 µm (Photometrics) and 488 nm, 561 nm, and 642 nm solid-state lasers (Coherent and MPB communications). Image stacks were acquired in 0.125 µm increments in the z-axis in sequential imaging mode. Samples were illuminated by a coherent scrambled laser light source first passed through a diffraction grating to generate the structured illumination by interference of light orders in the image plane to create a 3D sinusoidal pattern, with lateral stripes approximately 0.270 µm apart. The pattern was shifted laterally through five phases and through three angular rotations of 60° for each Z-section, separated by 0.125 µm. Exposure times were typically between 25 and 150 ms, and the power of each laser was adjusted to achieve optimal fluorescence intensities between 2,000 and 4,000 in a raw image of 16-bit dynamic range, at the lowest possible laser power to minimize photo bleaching. Color channels were carefully aligned using alignment parameter from control measurements with 0.5 µm diameter multi-spectral fluorescent beads (Thermo Fisher Scientific).

### Image analysis

All wide-field images presented in the figures were subjected to a constrained iterative deconvolution using softWoRx (GE Healthcare Life Sciences). All image analysis/quantification of fluorescence was performed using Fiji (Schindelin et al., 2012) on raw (i.e., nondeconvolved) images.

To quantify total fluorescence intensity of *α*-Nup188 and *α*-γ-Tubulin immunostaining at the centrosomes, cells were staged in mitosis by the morphology of the chromosomes stained with Hoechst. The integrated density of a region of interest that encompassed the centrosome was measured and divided by mean background (between cells) fluorescence.

To quantify SNAP-Nup188 fluorescence at centrosomes and the nuclear envelope during turnover experiments, a region of interest across the nuclear envelope and within the centrosome was drawn. The mean fluorescence intensity of these regions of interest were measured and subtracted from the mean fluorescence of intracellular cellular background on an individual cell basis.

To quantify changes in fluorescence intensity of *α*-PCM1 labeling around centrosomes, the radial profile ImageJ plugin was used. The total fluorescence intensity of an annulus of a given radius with the centrosome as its center was measured. To plot the cumulative distribution of *α*-PCM1 labeling throughout the cell, the radial fluorescence intensity distribution for each individual cell was normalized to the total fluorescence in each cell, averaged across all cells and normalized to the maximum value.

Centriole numbers Sas6 foci were quantified manually in each cell. To quantify changes in Sas6 levels at centrosomes, the mean fluorescence intensity of *α*-Sas6-foci centrosome was measured and the mean fluorescence the cellular background was subtracted.

The 3D-SIM images were subjected to SIM reconstruction and image processing using the SoftWoRx 3.7 imaging software package (GE Healthcare Life Sciences). The channels were then aligned in x, y, and rotationally using predetermined shifts as measured using a target lens and the SoftWoRx alignment tool (GE Healthcare Life Sciences).

To estimate the diameter of the ring distributions of PCM components surrounding centrioles, we fit experimental fluorescent images with a theoretical image of a uniformly fluorescent ring of a given radius *R* that would be generated for a given width *σ* of a point-spread function (PSF). The PSF was approximated as a 2-dimensional Gaussian. Consequently, the theoretical image *I(x,y)* was computed by integrating the Gaussian along a circular contour according Equation S1.

Equation S1: 

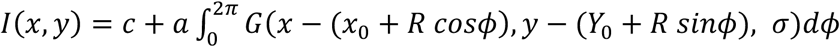

Here, *x_0_*, *y_0_* are coordinates of the ring center, *a* is a height (intensity) of the theoretical image, *c* is an image offset (i.e. background), and 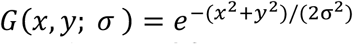. Equation S1 was numerically integrated using the Matlab built-in ‘integral’ function and then fitted to the experimental image using the Matlab ‘fit’ function. The following parameters were free to vary during the fitting: coordinates of the ring center *x_0_*, *y_0_*, ring radius *R*, width of PSF *σ*, image offset *c*, and image height *a*.

Input experimental images were chosen manually and cropped to regions that fully encompassed the structure of interest. These images were fitted automatically and used to generate plots and average images. To generate an average image of a structure formed by centrosome-associated proteins, all input images for fitting were aligned by using the fitting output (described above) to define a center of the structure and by cropping a square region of interest around the center. Subsequently, all independent images of the structure from different individual cells were averaged by calculating a mean pixel value for a given pixel position through all cropped images.

Note, that the average image (described above), while decreasing image noise, provides information at pixel resolution. Since the center of a ring structure can be at any position within the pixel, the experimental images of multiple independent rings contains information about the intensity distribution with sub-pixel resolution. To leverage this information, we calculated a radial distribution of intensity around the ring center. For each input image, distance from the ring center (defined by fitting) and the corresponding pixel intensity were calculated for all pixels in the image. These distance-intensity pairs were combined together and binned by the distance from the center to include *n* = 60 points in each bin. For each bin, a mean pixel value was calculated.

### Biotin Identification (BioID) proximity labeling

To determine the nearest neighbor interaction partners of BirA*-Nup188 and BirA* alone, we performed proximity-labeling using BioID (Roux et al., 2012). Following the protocol outlined in Firat-Karalar et al., (Firat-Karalar and Stearns, 2015) Biotin (Millipore Sigma) was added at a final concentration of 50 μM for 24 h to catalyze proximity-biotinylation. To affinity purify biotinylated proteins, cells were washed with PBS and then lysed in BioID lysis buffer (50 mM Tris, pH 7.4, 500 mM NaCl, 0.4% SDS, 5 mM EDTA, 1 mM DTT, 2% Triton X-100 and protease inhibitors (Thermo Fisher Scientific) for 10 min, on ice. To ensure lysis and to break-up chromatin, the mixtures were sonicated with a Bioruptor™ UCD-2000 (Diagenode). The resulting lysate was clarified by centrifugation at 14,000 rpm for 10 min at 4**°**C. The supernatant was collected and diluted with an equal volume of 50 mM Tris pH 7.4 to promote binding to streptavidin beads. Streptavidin-coupled magnetic beads (Dynabeads™ MyOne™ Streptavidin C1, Thermo Fisher Scientific) were added to the lysate and incubated at 4**°**C overnight. The beads were collected on a magnet and subjected to sequential washes with the following buffers: 1) 2% SDS, 2) 0.2% deoxycholate, 1% Triton X-100, 500 mM NaCl, 1 mM EDTA, 50 mM HEPES, pH 7.5, 3) 10 mM Tris, pH 8.1, 250 mM LiCl, 5% NP-40, 0.5% deoxycholate, 1% Triton X-100, 500 mM NaCl and 1 mM EDTA and 4) 50 mM Tris, pH 7.4, 50 mM NaCl.

### Mass Spectrometry

Affinity-purified biotinylated proteins were subjected to on-bead trypsin digestion. Briefly, beads were washed twice with 600 µl of 50 mM ammonium bicarbonate (pH 8.0) and then trypsin (Promega) was added (200 ng in 200 µl of 25 mM ammonium bicarbonate, pH 8) to carry out in-bead digestion for overnight at 37°C. After the supernatant was transferred to an Eppendorf tube, the beads were rinsed with 100 µl of water and the supernatants were combined. The solution was acidified by the adding 5 µl of 20% trifluoroacetic acid, and then dried in a SpeedVac (Thermo Fisher Scientific). Peptides were dissolved in 30 µl MS loading buffer (2% acetonitrile, 0.2% TFA), and protein concentration was determined by a Nanodrop™ (Thermo Fisher Scientific) A260/A280 measurement. An aliquot of each sample was then further diluted with loading buffer to 0.1 µg/µl and 5 µl (0.5 µg) was injected for LC-MS/MS analysis.

LC-MS/MS analysis was performed on a Thermo Scientific Orbitrap Elite mass spectrometer equipped with a Waters nanoAcquity UPLC system utilizing a binary solvent system (A: 100% water, 0.1% formic acid; B: 100% acetonitrile, 0.1% formic acid). Trapping was performed at 5 µl/min, 97% solvent A for 3 min using a Waters Symmetry® C18 180 µm x 20 mm trap column. Peptides were separated using an ACQUITY UPLC PST (BEH) C18 nanoACQUITY Column 1.7 µm, 75 µm x 250 mm (37°C) and eluted at 300 nl/min with the following gradient: 3% solvent B at initial conditions; 5% B at 3 min; 35% B at 140 min; 50% B at 155 min; 85% B at 160-165 min; return to initial conditions at 166 min. MS was acquired in the Orbitrap in profile mode over the 300-1,700 m/z range using 1 microscan, 30,000 resolution, AGC target of 1E6, and a full max ion time of 50 ms. Up to 15 MS/MS were collected per MS scan using collision induced dissociation (CID) on species with an intensity threshold of 5,000 and charge states 2 and above. Data dependent MS/MS were acquired in centroid mode in the ion trap using 1 microscan, AGC target of 2E4, full max IT of 100 ms, 2.0 m/z isolation window, and normalized collision energy of 35. Dynamic exclusion was enabled with a repeat count of 1, repeat duration of 30 sec, exclusion list size of 500, and exclusion duration of 60 sec.

All MS/MS spectra were searched in-house using the Mascot algorithm (version 2.6.0) for un-interpreted MS/MS spectra after using the Mascot Distiller program to generate Mascot compatible files. The data was searched against the Swiss Protein database with taxonomy restricted to *Homo sapiens*, allowing for methionine oxidation as a variable modification. Peptide mass tolerance was set to 10 ppm and MS/MS fragment tolerance to 0.5 Da. Normal and decoy database searches were run to determine the false discovery rates, and the confidence level was set to 95% within the MASCOT search engine for protein hits based on randomness.

Scaffold (version Scaffold_4.8.2, Proteome Software Inc.) was used to validate MS/MS based peptide and protein identifications. Peptide identifications were accepted if they could be established at greater than 95.0% probability by the Scaffold Local FDR algorithm. Protein identifications were accepted if they could be established at greater than 95.0% probability and contained at least 1 identified peptide.

To compare data across different runs and assess the abundance of each proximity partner, normalized spectral counts were determined for BirA* and BirA*-Nup188 samples. A spectral abundance factor (SAF) was calculated by normalizing the total spectral counts for a given protein to its length. To account for variability between independent runs, individual SAF values were then normalized against the sum of all SAFs for a particular run, resulting in a normalized SAF (NSAF). Only proteins that were 1) specific to Bir*-Nup188 2) had NSAF values 2.5 fold greater than found in the BirA-control and 3) were detected in at least 2 biological replicates were selected.

Gene ontology (GO) enrichment analysis for biological process was performed on the ranked list of NSAF values using GOrilla (Eden et al., 2009) followed by GO term redundancy reduction performed by REVIGO (Supek et al., 2011).

### Statistical analysis

All experiments were carried out at least three times. Graphs and statistical analyses were generated using Prism (GraphPad 7.0). *P*-values in all graphs were generated with tests as indicated in figure legends and are represented as follows: ns, *P* >0.05; * *P* ≤0.05; ** *P* ≤0.01 *** *P* ≤0.001, **** *P* ≤0.0001. All error bars represent the standard deviation from the mean.

## Acknowledgements

We thank Jean Kanyo at the Yale Keck Biotechnology Resource Laboratory for help with mass spectrometry analysis and Felix E. Rivera-Molina for expertise with 3D-SIM. This work was generously supported by grants from the National Institutes of Health (NIH): RO1 HL124402 to C.P.L and M.K.K, which supports N.V. and K.D. M.C. is funded by NIH-5F31HL134272 and NIH-5T32GM007223.

## Figure Legends

**Figure S1.**
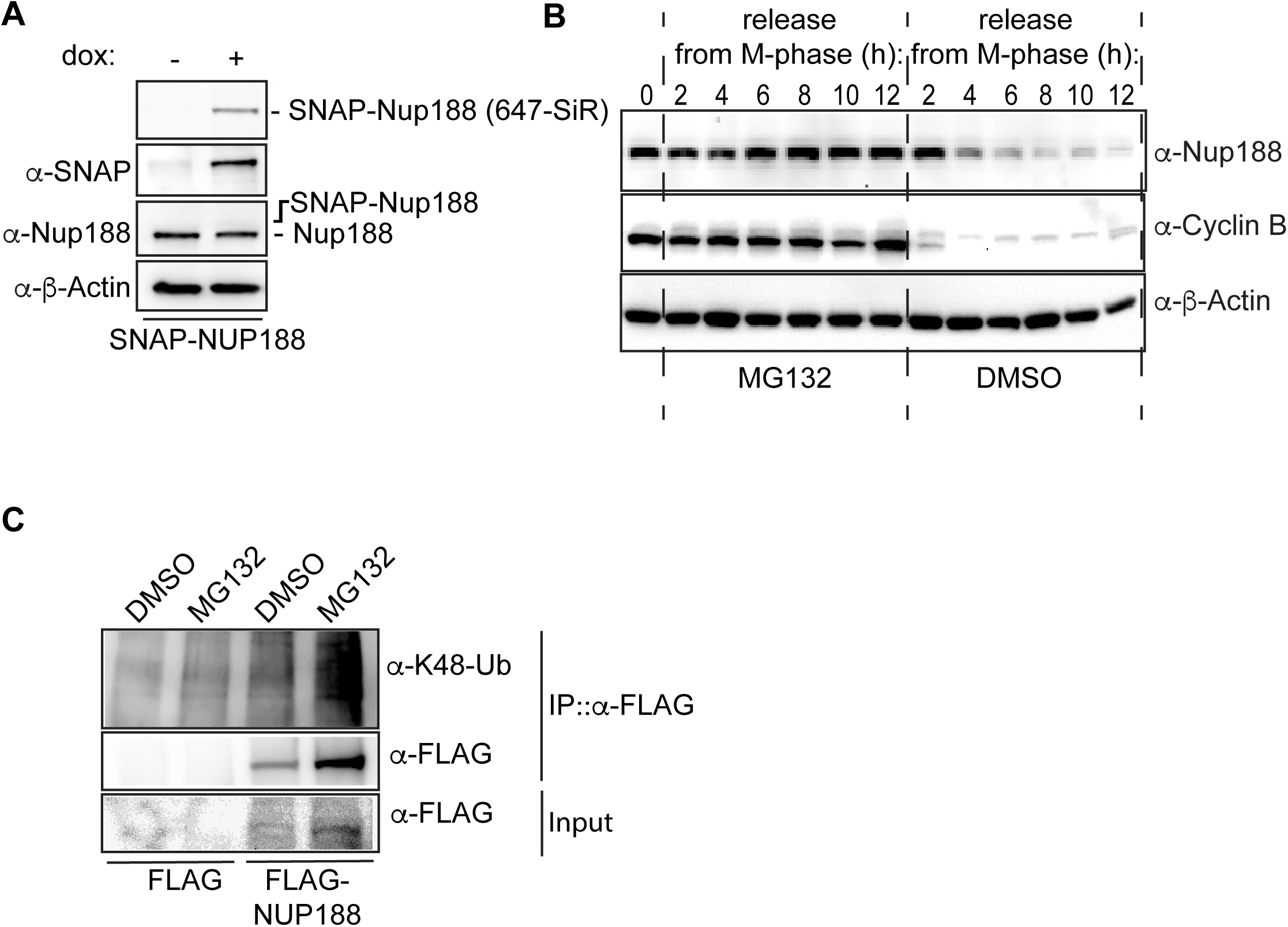
Nup188 is stabilized upon proteasome-inhibition. **(A)** SNAP-Nup188 levels are below endogenous Nup188. Western blot (and fluorescence labeling of SNAP-Nup188 with a 647-SiR dye, top panel) of total protein extracts derived from SNAP-NUP188-expressing HeLa cells collected after 48 h in the presence (+) or absence (-) of dox. Note that the putative position of SNAP-Nup188 is indicated as it is expressed at levels that cannot be detected with the *α*-Nup188 antibody. *α*-β-Actin is a protein load reference. **(B)** Inhibition of the proteasome stabilizes Nup188 levels leaving mitosis. Nocodazole-arrested HeLa-M cells were released in the presence of MG132 or carrier alone (DMSO). Western blots of total protein extracts were performed at the indicated time points. *α*-β-Actin is a protein load reference. **(C)** Ubiquitylated species bound to Nup188 are enriched upon proteasome inhibition. Western blots of input (2%) and immunoprecipitated (IP) fractions (100%) from IPs of FLAG-Nup188 or FLAG-alone from cell extracts treated with MG132 or DMSO. FLAG-Nup188 was detected with an *α*-Flag antibody and ubiquitylation using *α*-K48-Ub antibody.

**Figure S2.**
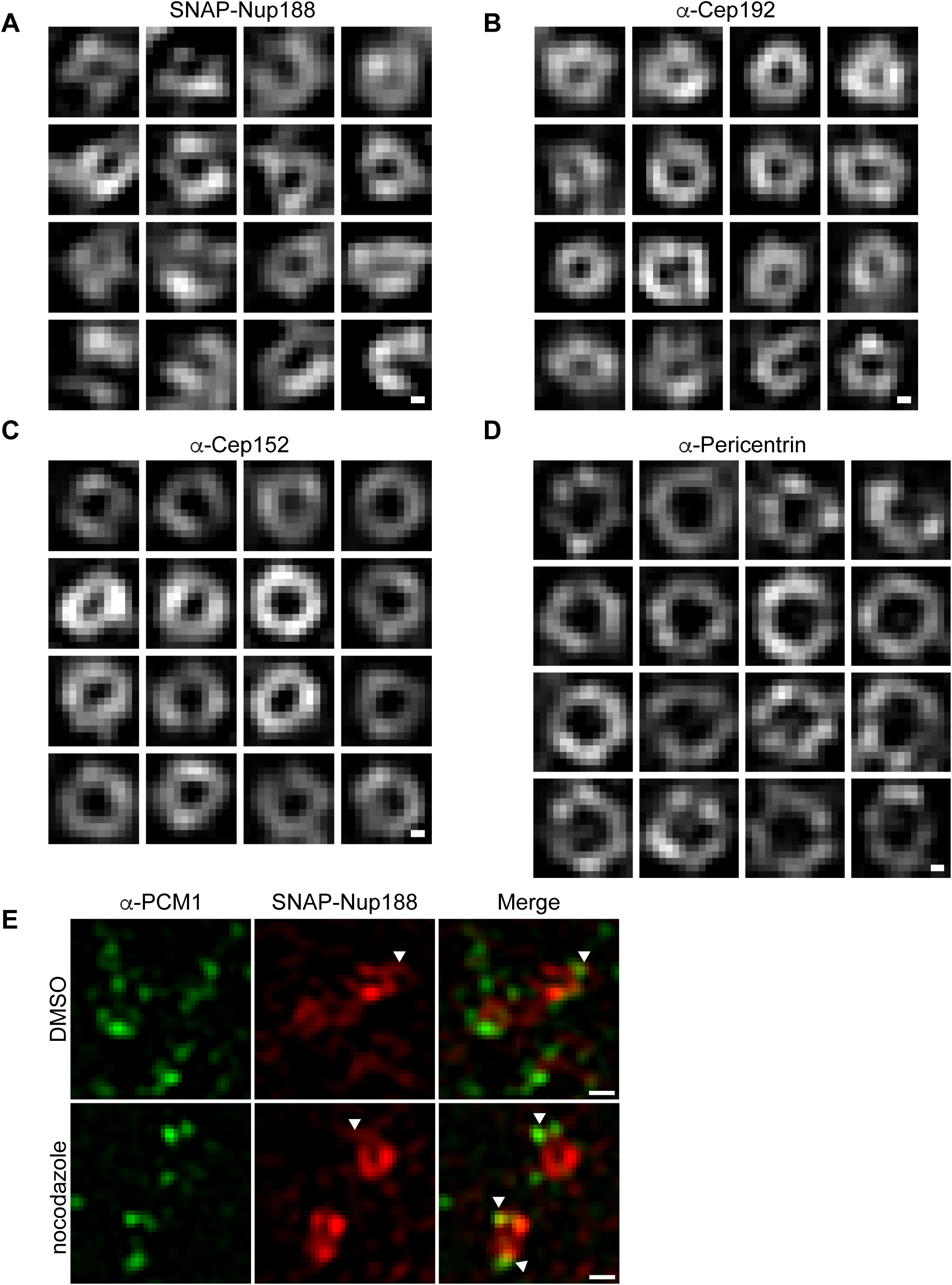
Nup188 centrosome labeling has extensions that emanate from its circular core. **(A-D)** 3D-SIM image montages of centrosomes labeled with TMR-Star (SNAP-Nup188) and indicated centrosome proteins. Scale bar is 0.10 µm. **(E)** Neither Nup188 extensions, nor *α*-PCM1 labeling at centrosomes are perturbed by microtubule disruption. 3D-SIM images of SNAP-Nup188 (TMR-Star dye, red) and *α*-PCM1 (green) at centrosomes in cells treated with nocodazole or DMSO. Arrowheads point to extensions of SNAP-Nup188, which sometimes co-localize with the *α*-PCM1 label. Scale bar, 0.25 µm.

**Figure S3.**
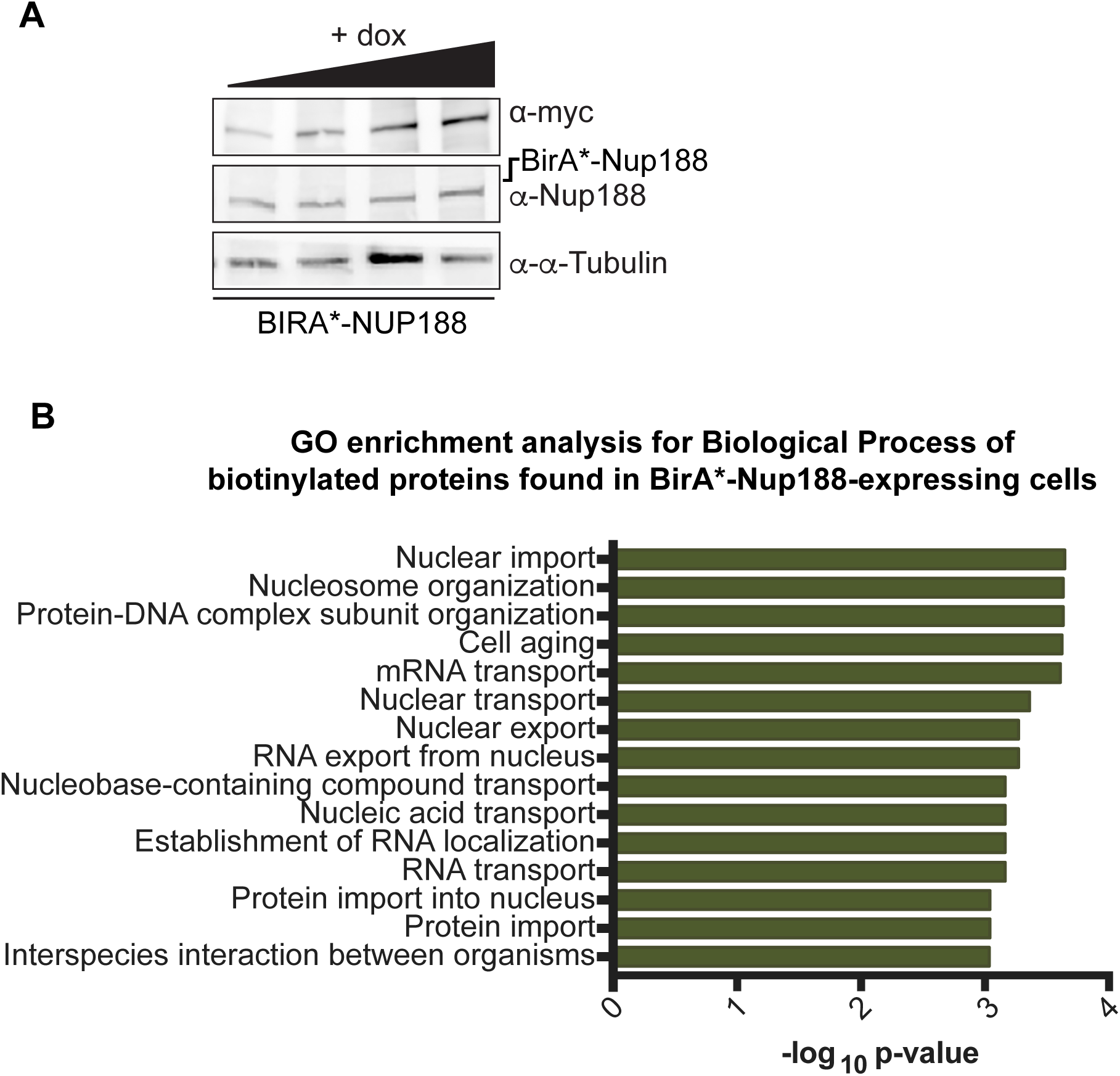
BirA*-Nup188 is produced below endogenous Nup188 levels and leads to biotinylation of nuclear-function proteins. **(A)**. Western blots examining the levels of BirA*-Nup188 in total protein samples derived from SNAP-NUP188-expressing HeLa cells treated with increasing amounts of dox. Expected position of BirA*-Nup188 is indicated relative to the observed endogenous Nup188; *α*-*α*-Tubulin is a reference for total protein loads. **(B)** BirA*-Nup188 proximity-dependent biotin labeling identifies nuclear components. Plot showing Gene Ontology (GO) enrichment analysis for biological process performed on the ranked list of NSAF values in Table S1. GOrilla (Eden et al., 2009) followed by GO term redundancy reduction by REVIGO (Supek et al., 2011) was performed.

**Figure S4.**
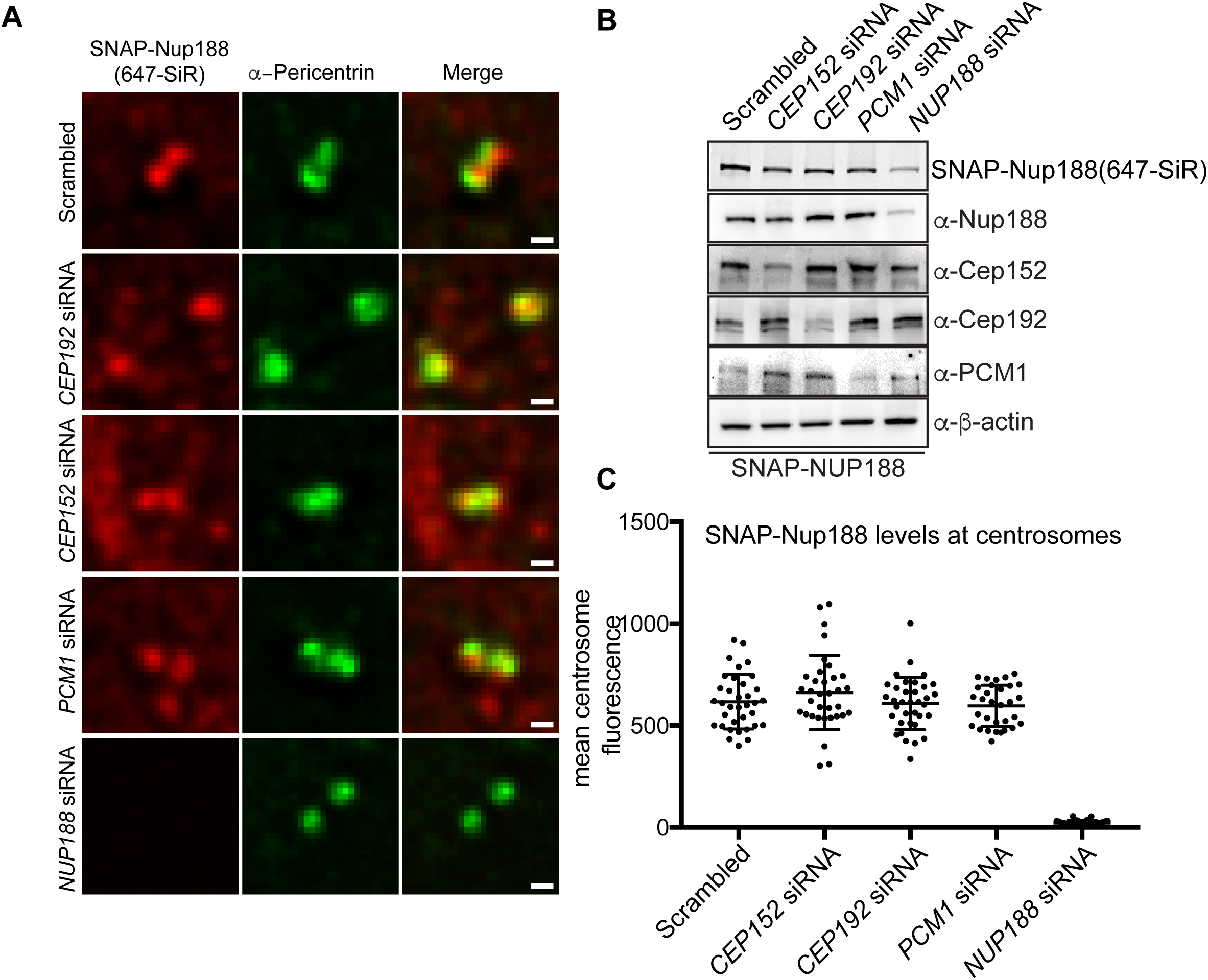
Knockdown of PCM components does not affect SNAP-Nup188 loading at centrosomes. **(A)** Fluorescence micrographs of SNAP-Nup188 (labeled with 647-SiR dye/red) and *α*-Pericentrin (green) in HeLa cells expressing SNAP-Nup188, treated with the indicated siRNAs. **(B)** Western blots (or fluorescence image/top panel) of total protein from SNAP-NUP188-expressing HeLa cell extracts prepared from cells treated with the indicated siRNAs. *α*-β-Actin is a protein load reference. **(C)** Plot of the mean fluorescence at centrosomes of SNAP-Nup188 labeled with 647-SiR dye in cells treated with indicated siRNAs. >30 cells were quantified with mean ±SD shown.

**Figure S5.**
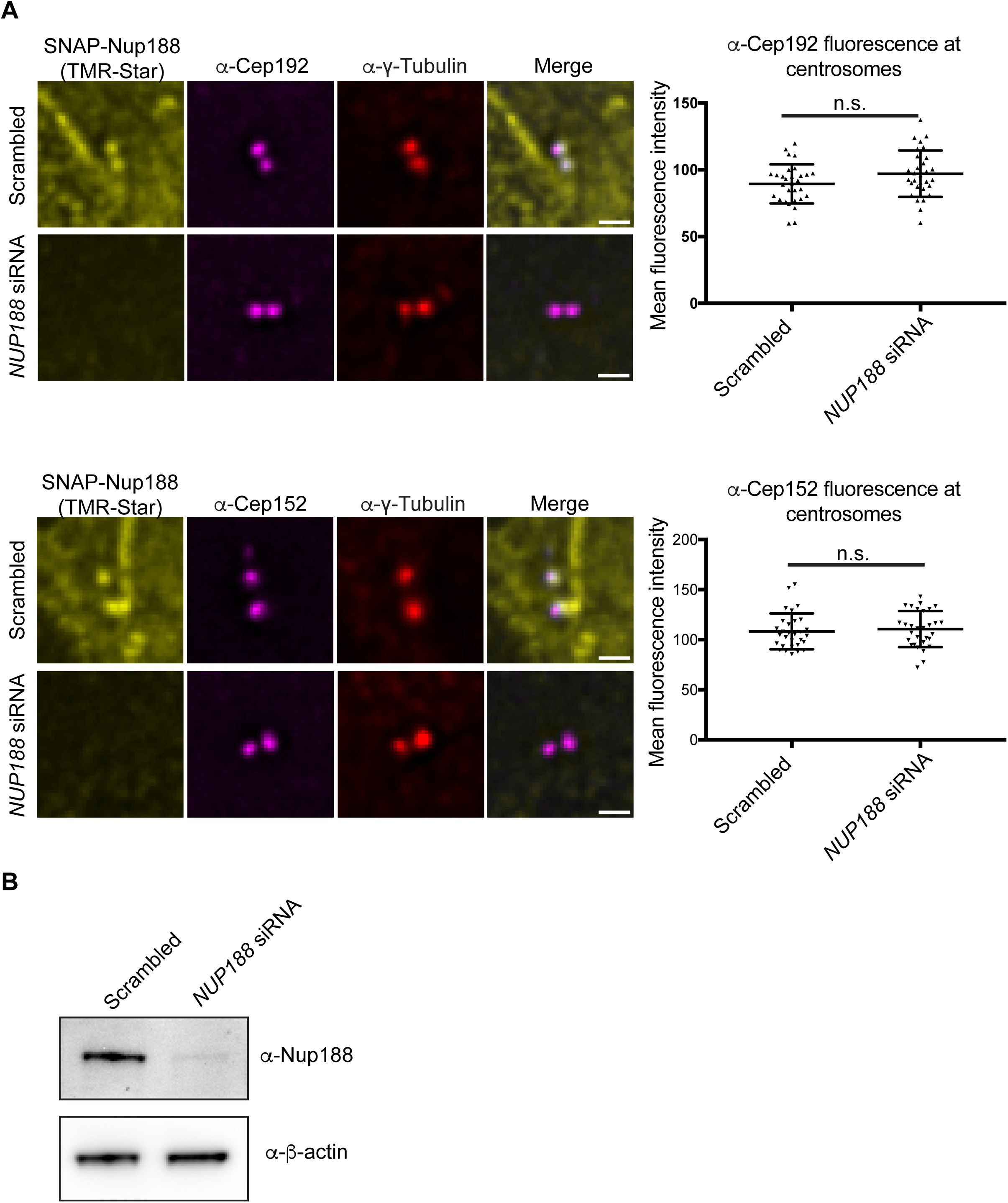
Nup188 depletion does not impact Cep192 and Cep152 association with centrosomes. **(A)** Fluorescence micrographs of HeLa cells expressing SNAP-Nup188 (TMR-Star) treated with *NUP188* or scrambled siRNAs and labeled with *α*-Cep192 (magenta, top), *α*-Cep152 (magenta, bottom) and *α*-γ-Tubulin (red). Scale bar, 1 µm. Representative plot of the mean fluorescence +/− SD at centrosomes (n>30). Error bars indicate ±SD. n.s. is not significant as calculated from two-way ANOVA. **(B)** Western blot of total protein from cells treated with scrambled or *NUP188* siRNA. *α*-β-Actin is a protein load reference.

**Figure S6.**
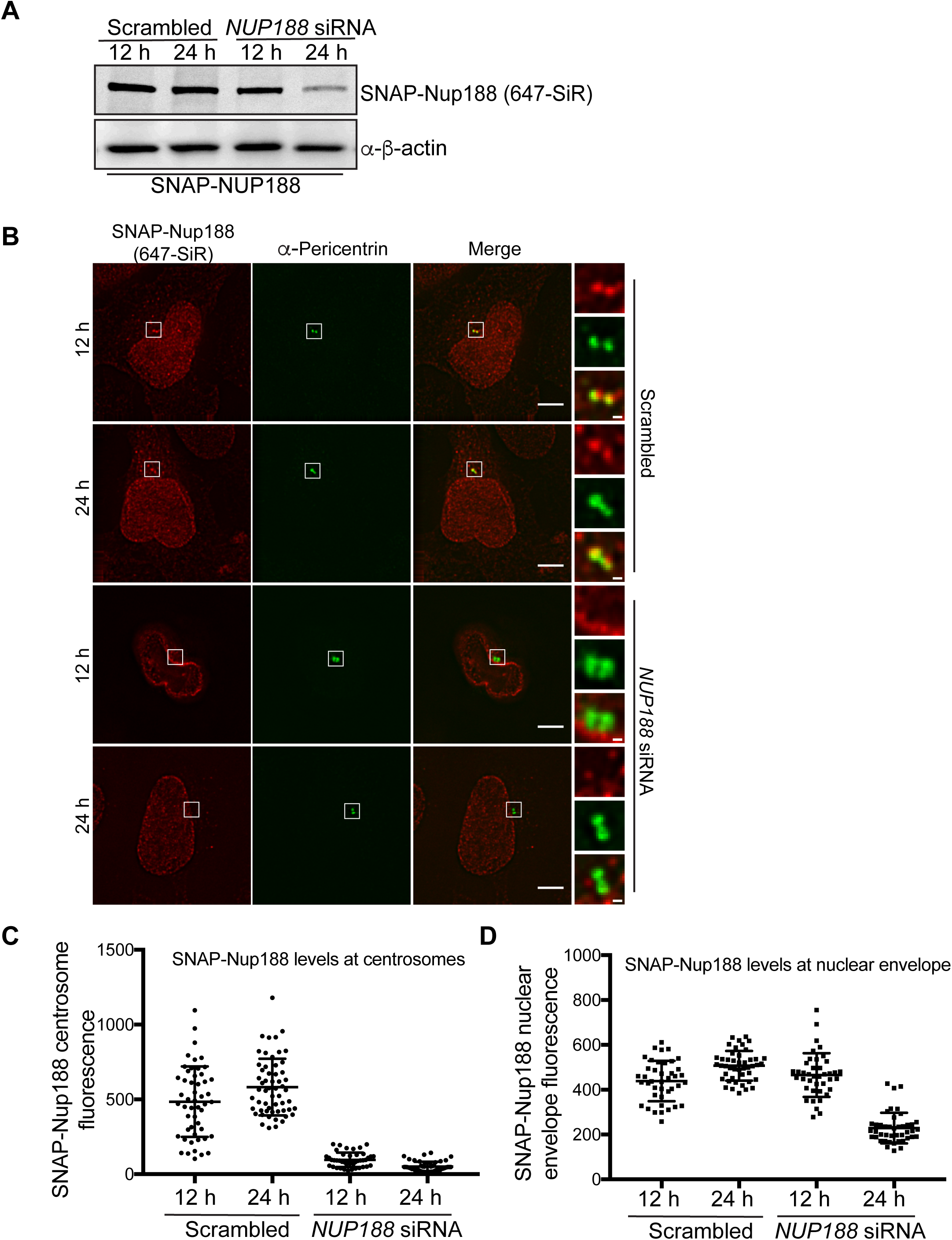
The centrosomal pool of Nup188 is sensitive to siRNA knockdown. **(A)** Western blots (or fluorescence image/top panel) of total protein from SNAP-NUP188-expressing HeLa cell extracts prepared from cells treated with the indicated siRNAs for 12 or 24 h. *α*-β-Actin is a protein load reference. **(B)** Fluorescence micrographs of SNAP-Nup188 (labeled with 647-SiR/red) in HeLa cells treated with *NUP188* or scrambled siRNA for 12 or 24 h. Centrosomes are labeled with *α*-Pericentrin (green). Scale bar, 5 μm. Magnifications of boxed region are shown at right. Scale bar, 0.5 μm. **(C, D)** Plots of mean SNAP-Nup188 fluorescence (647-SiR) +/− SD at centrosomes and the nuclear envelope in the indicated conditions where >30 cells were quantified.

**Figure S7.**
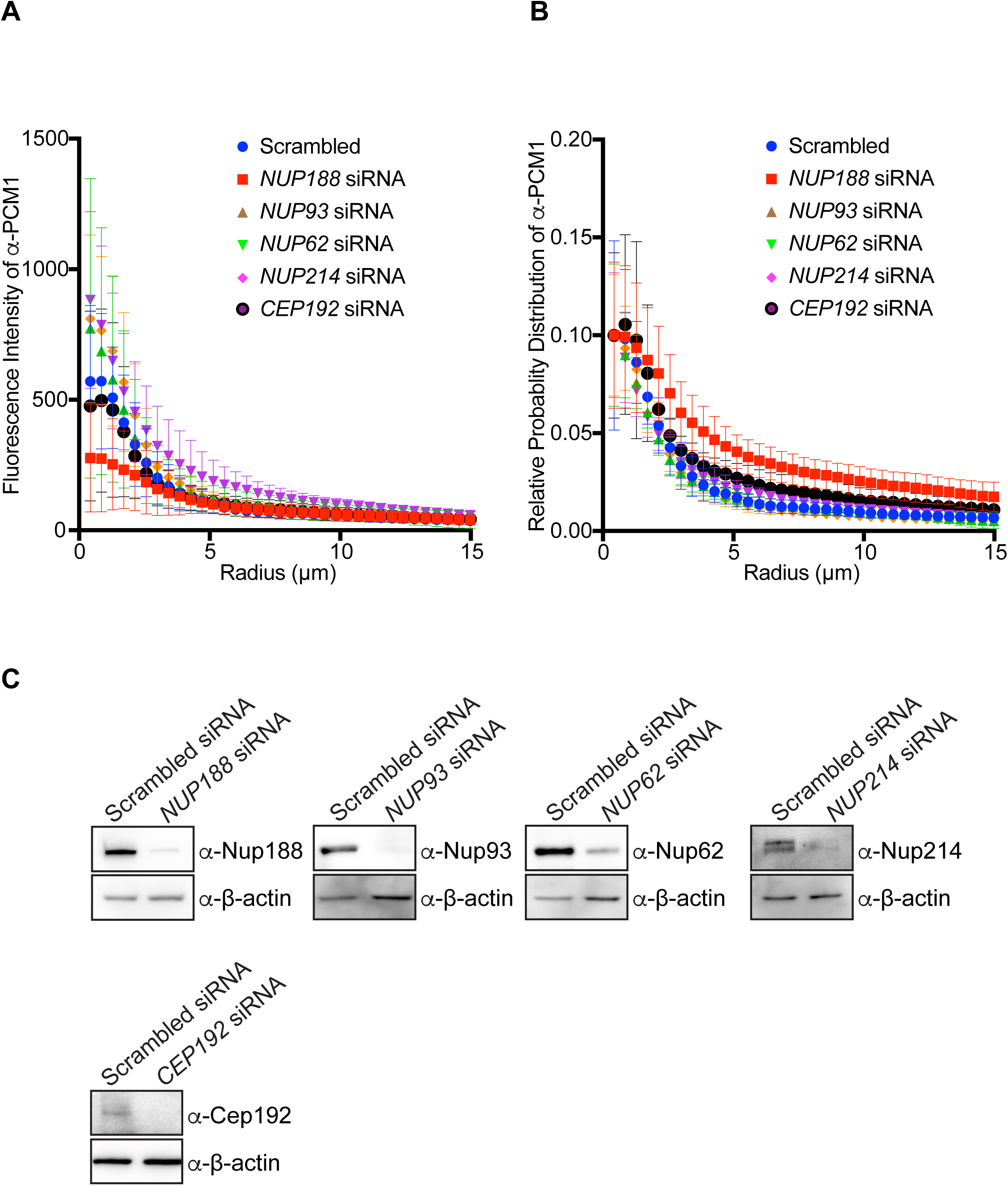
Knockdown of Nup188 specifically impacts centriolar satellite distribution. **(A)** Plot of the total fluorescence of *α*-PCM1 labeled centriolar satellites within an annulus of indicated radii from the center of *α*-Centrin fluorescence (n=10, see images in Figure 6 for example). Error bars indicate ±SD. **(B)** As in (A) but plotting the cumulative distribution of *α*-PCM1 labeling throughout the cell (n=10). Error bars indicate ±SD. **(C)** Western blots of total protein from HeLa cell extracts treated with the indicated siRNA. *α*-β-Actin is a protein load reference.

**Figure S8.**
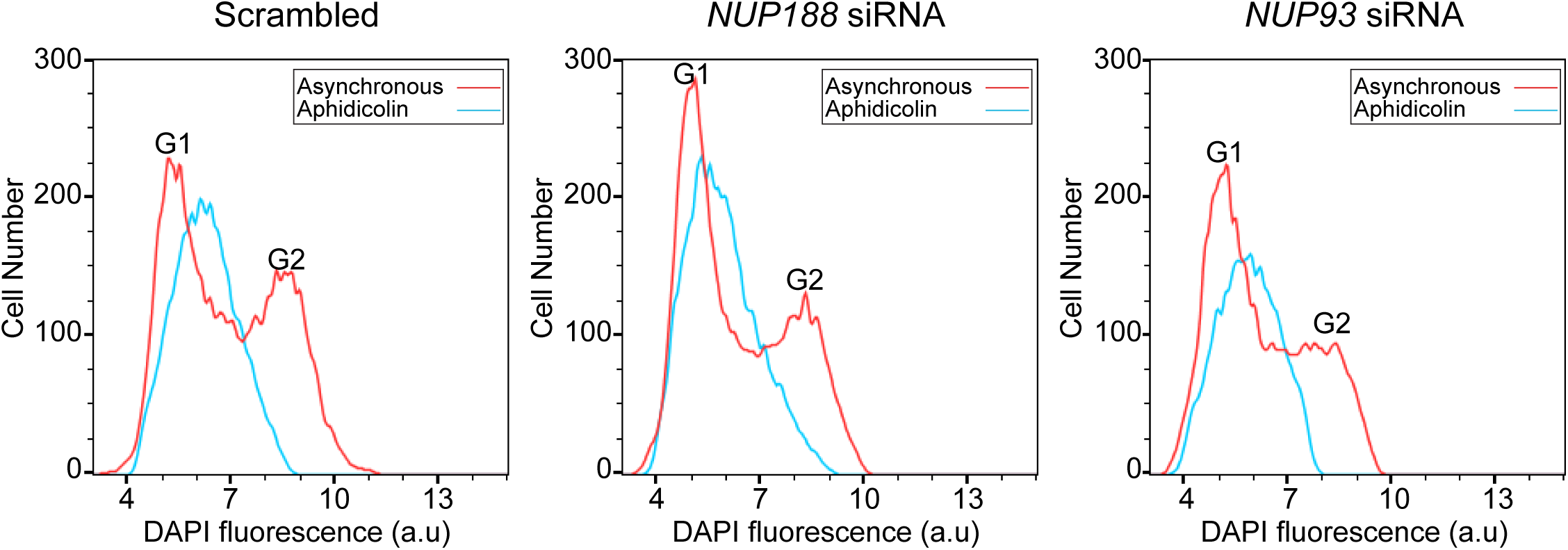
FACS analysis confirms S-phase arrest in *NUP188* siRNA treated cells. FACS profiles of DNA content (from DAPI fluorescence in arbitrary units (a.u)) in asynchronous (red lines) and S-phase synchronized (aphidicolin, blue lines) U2OS cells treated with the indicated siRNAs.

---

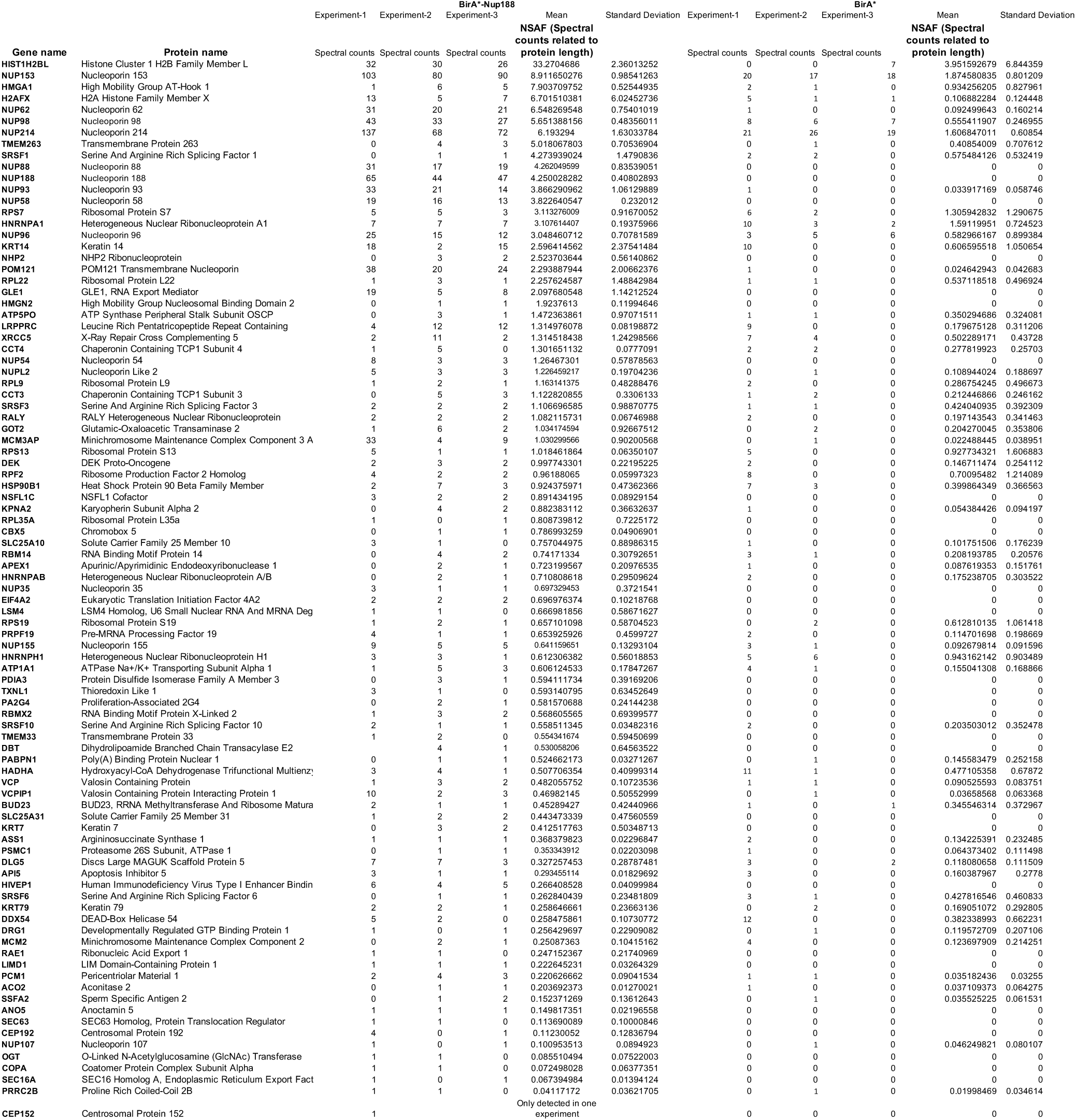

**Table S2:**
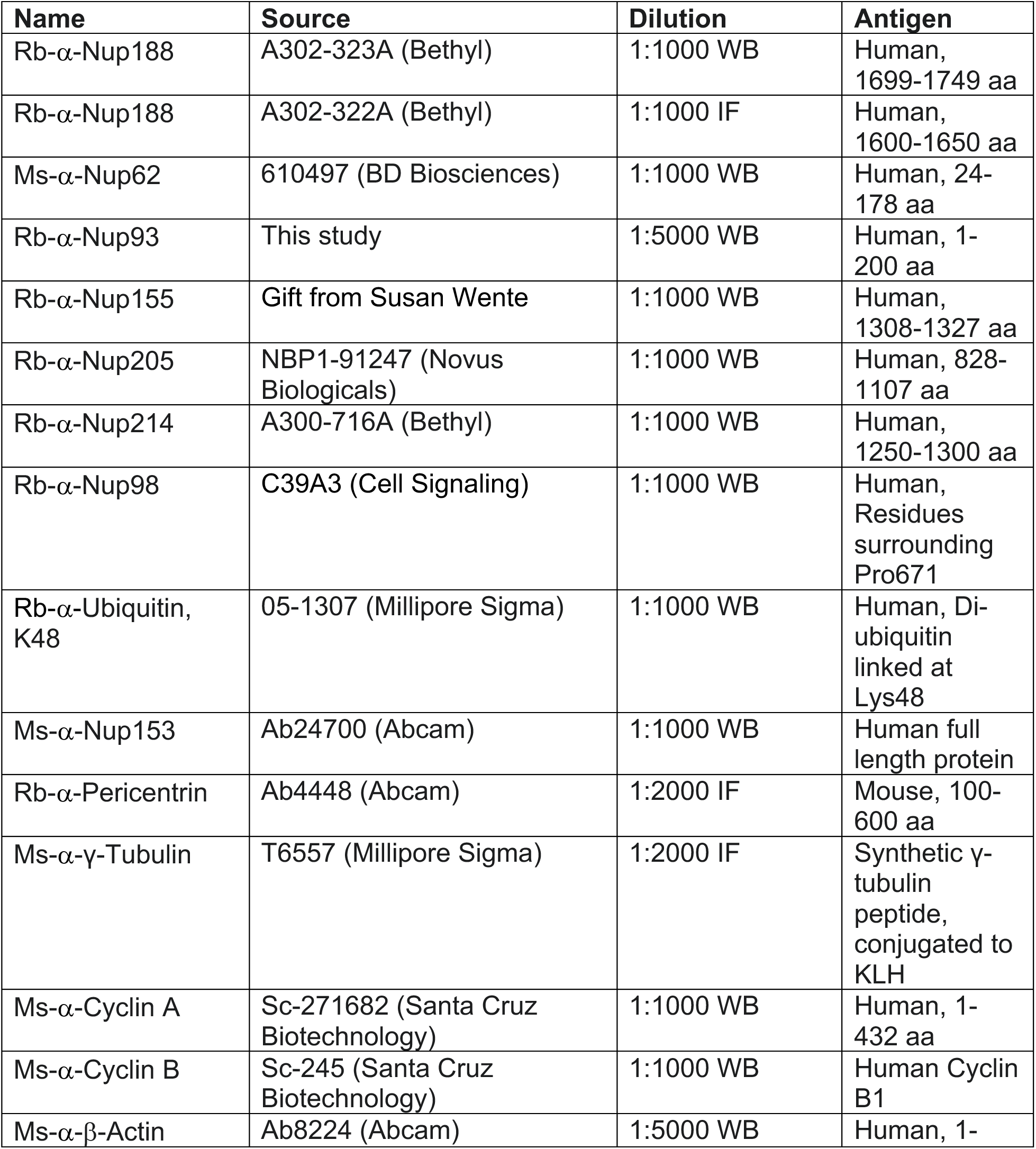
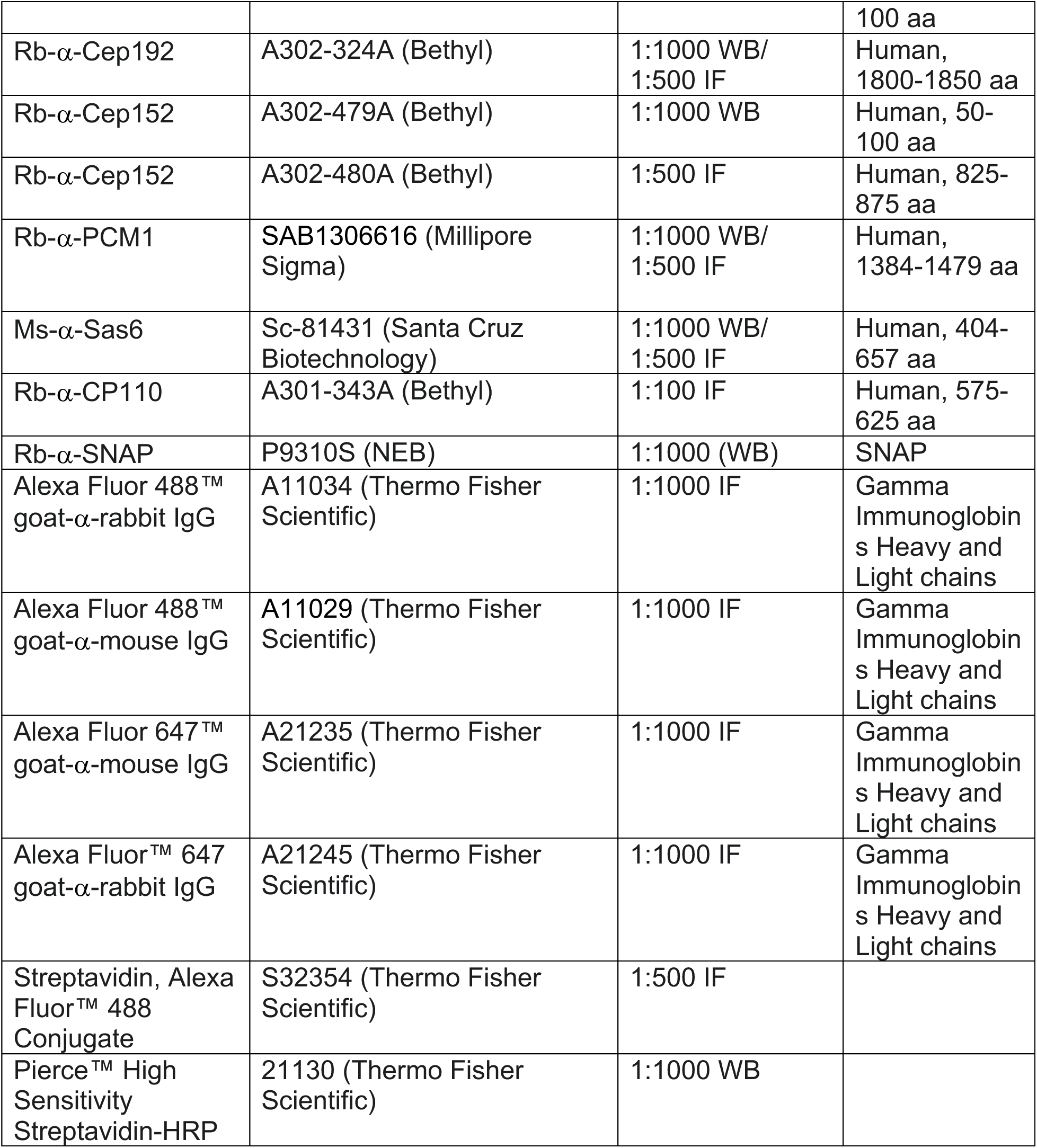
Antibodies and Conjugated Streptavidin.

| **Name** | **Source** | **Dilution** | **Antigen** |
| --- | --- | --- | --- |
| Rb-α-Nup188 | A302-323A (Bethyl) | 1:1000 WB | Human, 1699-1749 aa |
| Rb-α-Nup188 | A302-322A (Bethyl) | 1:1000 IF | Human, 1600-1650 aa |
| Ms-α-Nup62 | 610497 (BD Biosciences) | 1:1000 WB | Human, 24-178 aa |
| Rb-α-Nup93 | This study | 1:5000 WB | Human, 1-200 aa |
| Rb-α-Nup155 | Gift from Susan Wente | 1:1000 WB | Human, 1308-1327 aa |
| Rb-α-Nup205 | NBP1-91247 (Novus Biologicals) | 1:1000 WB | Human, 828-1107 aa |
| Rb-α-Nup214 | A300-716A (Bethyl) | 1:1000 WB | Human, 1250-1300 aa |
| Rb-α-Nup98 | C39A3 (Cell Signaling) | 1:1000 WB | Human, Residues surrounding Pro671 |
| Rb-α-Ubiquitin, K48 | 05-1307 (Millipore Sigma) | 1:1000 WB | Human, Di-ubiquitin linked at Lys48 |
| Ms-α-Nup153 | Ab24700 (Abcam) | 1:1000 WB | Human full length protein |
| Rb-α-Pericentrin | Ab4448 (Abcam) | 1:2000 IF | Mouse, 100-600 aa |
| Ms-α-γ-Tubulin | T6557 (Millipore Sigma) | 1:2000 IF | Synthetic γ-tubulin peptide, conjugated to KLH |
| Ms-α-Cyclin A | Sc-271682 (Santa Cruz Biotechnology) | 1:1000 WB | Human, 1-432 aa |
| Ms-α-Cyclin B | Sc-245 (Santa Cruz Biotechnology) | 1:1000 WB | Human Cyclin B1 |
| Ms-α-β-Actin | Ab8224 (Abcam) | 1:5000 WB | Human, 1- |
|  |  |  | 100 aa |
| Rb-α-Cep192 | A302-324A (Bethyl) | 1:1000 WB/<br>1:500 IF | Human,<br>1800-1850 aa |
| Rb-α-Cep152 | A302-479A (Bethyl) | 1:1000 WB | Human, 50-100 aa |
| Rb-α-Cep152 | A302-480A (Bethyl) | 1:500 IF | Human, 825-875 aa |
| Rb-α-PCM1 | SAB1306616 (Millipore<br>Sigma) | 1:1000 WB/<br>1:500 IF | Human,<br>1384-1479 aa |
| Ms-α-Sas6 | Sc-81431 (Santa Cruz<br>Biotechnology) | 1:1000 WB/<br>1:500 IF | Human, 404-657 aa |
| Rb-α-CP110 | A301-343A (Bethyl) | 1:100 IF | Human, 575-625 aa |
| Rb-α-SNAP | P9310S (NEB) | 1:1000 (WB) | SNAP |
| Alexa Fluor 488<sup>TM</sup><br>goat-α-rabbit IgG | A11034 (Thermo Fisher<br>Scientific) | 1:1000 IF | Gamma<br>Immunoglobulin<br>s Heavy and<br>Light chains |
| Alexa Fluor 488<sup>TM</sup><br>goat-α-mouse IgG | A11029 (Thermo Fisher<br>Scientific) | 1:1000 IF | Gamma<br>Immunoglobulin<br>s Heavy and<br>Light chains |
| Alexa Fluor 647<sup>TM</sup><br>goat-α-mouse IgG | A21235 (Thermo Fisher<br>Scientific) | 1:1000 IF | Gamma<br>Immunoglobulin<br>s Heavy and<br>Light chains |
| Alexa Fluor<sup>TM</sup> 647<br>goat-α-rabbit IgG | A21245 (Thermo Fisher<br>Scientific) | 1:1000 IF | Gamma<br>Immunoglobulin<br>s Heavy and<br>Light chains |
| Streptavidin, Alexa<br>Fluor<sup>TM</sup> 488<br>Conjugate | S32354 (Thermo Fisher<br>Scientific) | 1:500 IF |  |
| Pierce<sup>TM</sup> High<br>Sensitivity<br>Streptavidin-HRP | 21130 (Thermo Fisher<br>Scientific) | 1:1000 WB |  |

**Table S3:**
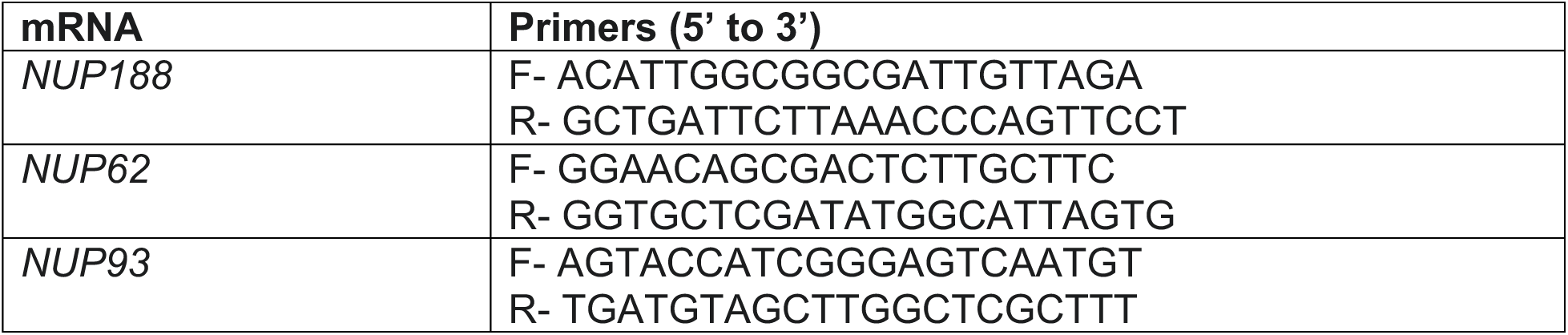
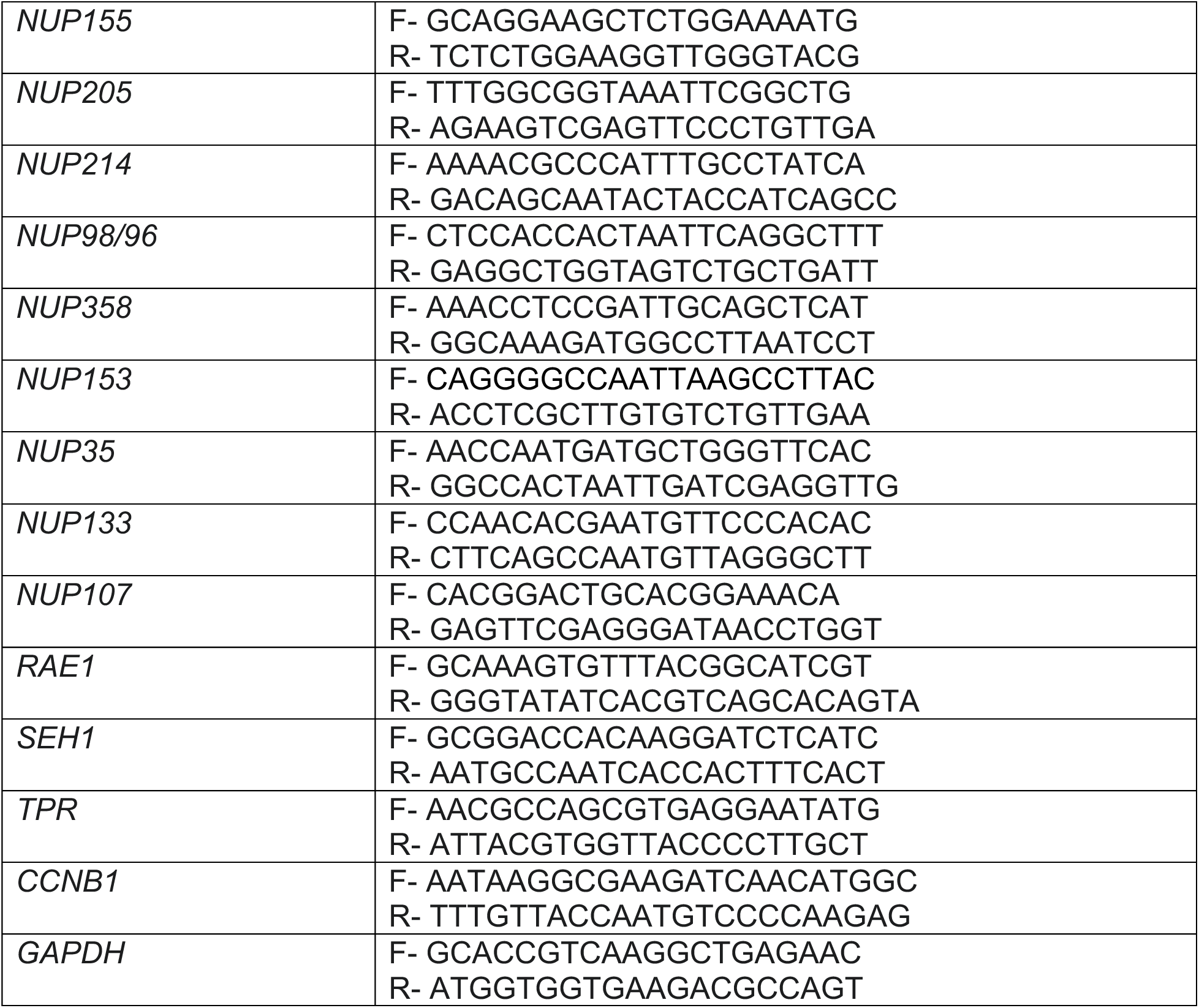
RT-qPCR Primers.

**Table S4:**
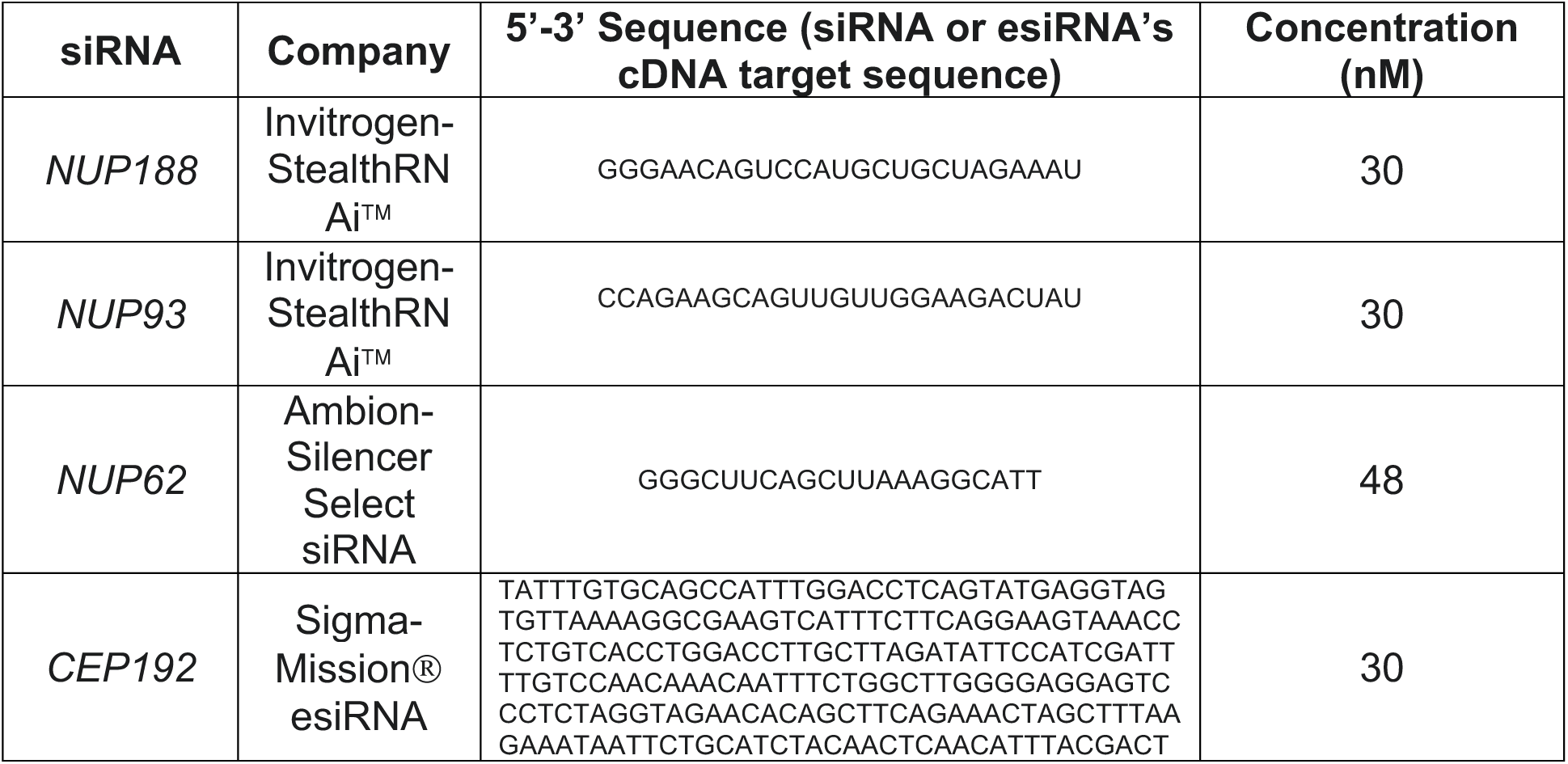
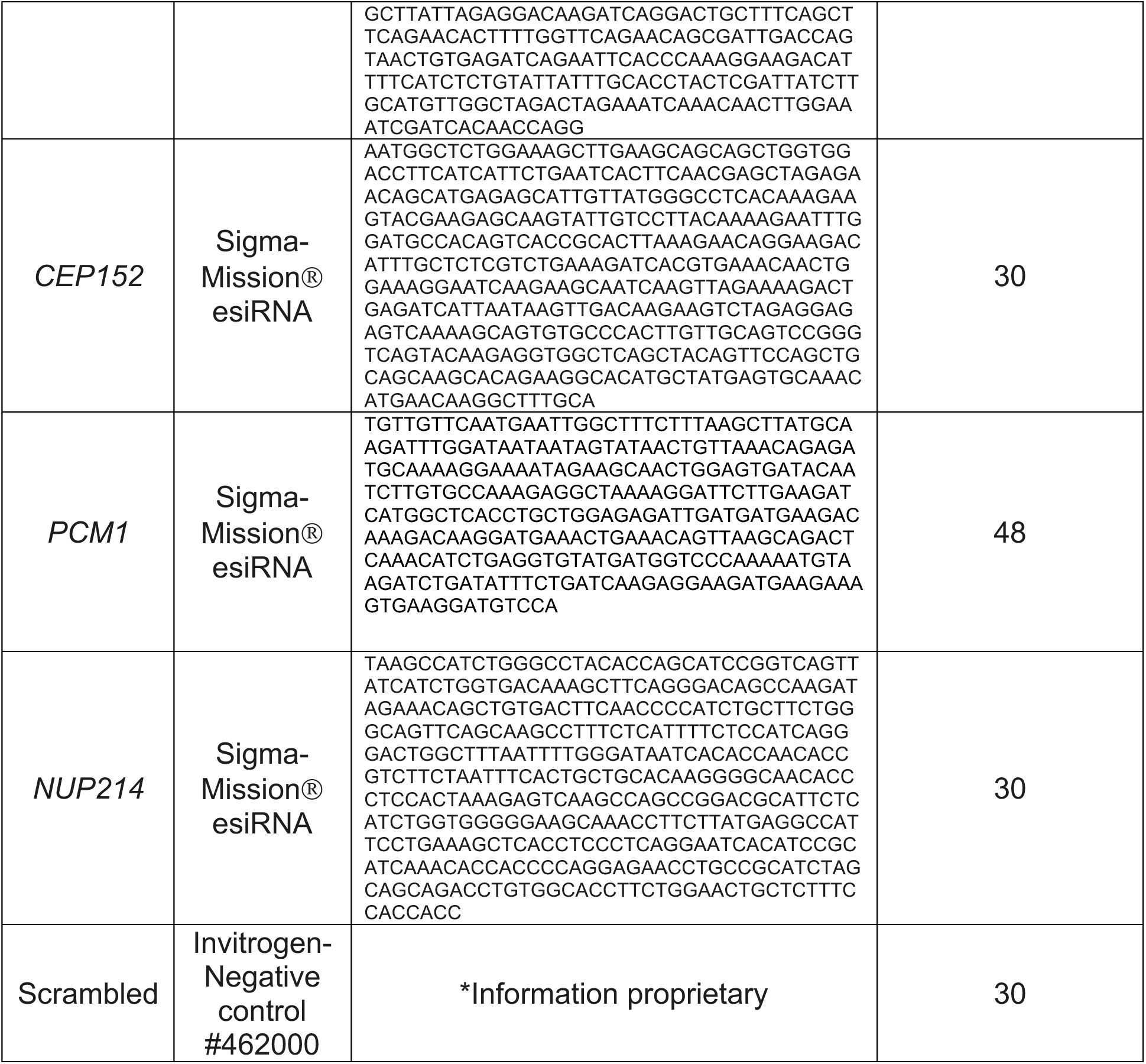
siRNA.

**Table S5:** Plasmids.

| Plasmid | Source | Description |
| --- | --- | --- |
| pNV1 | This study | BIRA* in pcDNA5/FRT/TO |
| pNV2 | This study | SNAP-NUP188 in pcDNA5/FRT/TO |
| pNV3 | This study | BIRA*-NUP188 in pcDNA5/FRT/TO |
| pMC1 | This study | SNAP replacing GFP in GFP-NUP188 |
| pMC2 | This study | BIRA* replacing GFP in GFP-NUP188 |
| pKD1 | This study | FLAG-NUP188 in p3XFLAG-CMV™-10 |
| p3XFLAG-CMV™-10 | Millipore-Sigma<br>(#E7658) | Vector for expression of N-terminally<br>3xFLAG-tagged proteins in mammalian<br>cells |
| pOG44 | Thermo Fisher<br>Scientific<br>(#V600520) | Flp-Recombinase Expression Vector |
| pcDNA5/FRT/TO | Thermo Fisher Scientific (#V652020) | Genomic integration vector for tetracycline-inducible expression of proteins in mammalian cells. |
| pSNAPf | NEB (#N9183) | Vector for expression of SNAP-tagged proteins in mammalian cells |
| pcDNA3.1 mycBioID | Addgene (#35700) | Vector for expression of BIRA*-tagged proteins in mammalian cells for BioID assay |
| GFP-NUP188 | Gift from Dr. K Tanaka | GFP-NUP188 in pEGFP-C1 |
| FLAG-NUP188 | Gift from Dr. K Tanaka | FLAG-NUP188 in pcDNA5/FRT |

